# Inferring Ancestry with the Hierarchical Soft Clustering Approach tangleGen

**DOI:** 10.1101/2024.03.27.586940

**Authors:** Klara Elisabeth Burger, Solveig Klepper, Ulrike von Luxburg, Franz Baumdicker

**Affiliations:** Cluster of Excellence “Machine Learning: New Perspectives for Science”, University of Tübingen, Germany; Department of Computer Science, University of Tübingen, Germany; Tübingen AI Center; Cluster of Excellence “Controlling Microbes to Fight Infections”, Mathematical and Computational Population Genetics, University of Tübingen, Germany; Institute for Bioinformatics and Medical Informatics (IBMI), University of Tübingen, Germany

**Keywords:** Tangles, ADMIXTURE, ancestry inference, hierarchical clustering, population genetics

## Abstract

Understanding the genetic ancestry of populations is central to numerous scientific and societal fields. It contributes to a better understanding of human evolutionary history, advances personalized medicine, aids in forensic identification, and allows individuals to connect to their genealogical roots. Existing methods, such as ADMIXTURE, have significantly improved our ability to infer ancestries. However, these methods typically work with a fixed number of independent ancestral populations. As a result, they provide insight into genetic admixture, but do not include a hierarchical interpretation. In particular, the intricate ancestral population structures remain difficult to unravel. Alternative methods with a consistent inheritance structure, such as hierarchical clustering, may offer benefits in terms of interpreting the inferred ancestries. Here, we present tangleGen, a soft clustering tool that transfers the hierarchical machine learning framework Tangles, which leverages graph theoretical concepts, to the field of population genetics. The hierarchical perspective of tangleGen on the composition and structure of populations improves the interpretability of the inferred ancestral relationships. Moreover, tangleGen adds a new layer of explainability, as it allows identifying the SNPs that are responsible for the clustering structure. We demonstrate the capabilities and benefits of tangleGen for the inference of ancestral relationships, using both simulated data and data from the 1000 Genomes Project.

## Introduction

Inference of population structure is central to many studies (Mathieson and Scally, 2020; Padhukasahasram, 2014). Insights into the genetic ancestry of populations contribute to a better understanding of human evolutionary history (Reich et al., 2009) and advances personalized medicine (Rotimi and Jorde, 2010; Reich et al., 2009), forensic identification (Pfaffelhuber et al., 2022), but also the conservation of endangered species (Pearse and Crandall, 2004; Randi, 2008). The analysis relies on genomic data, specifically the sequencing of single nucleotide polymorphisms (SNPs) for each individual. Distinct populations exhibit varying allele frequencies for specific SNPs, which allows conclusions to be drawn about the population structure. This information is often used as a basis to infer ancestries.

The development of methods to infer population structure has been part of research for decades, with both model-based (Pritchard et al., 2000; Tang et al., 2005; Raj et al., 2014; Behr et al., 2016; Wang, 2022; Dominguez Mantes et al., 2023; Chiu et al., 2022) and model-free approaches (Patterson et al., 2006; Jombart et al., 2010) being pursued. For a compact summary, we recommend Wang (2022). A popular modeling framework was introduced by Pritchard et al. (2000) in a Bayesian setting and also serves as the basis of ADMIXTURE by Alexander et al. (2009), a maximum likelihood method for the simultaneous estimation of ancestry proportions of individuals and allele frequencies in populations. The probabilistic model is based on biallelic SNP genotypes, where SNPs are assumed to be independent and originate from a fixed number of independent ancestral populations. The inferred ancestry proportions are often difficult to interpret due to the model assumptions, inference methods, and the complexity of actual demographic histories (Lawson et al., 2018). For example, in ADMIXTURE the user has to pre-specify the number of ancestral populations, which influences the inferred ancestry proportions. Since this number is typically unknown, it must be estimated (Wang, 2022; Lawson et al., 2018). In practice, results are usually considered for a range of numbers of ancestral populations. However, results can provide inconsistent predictions for different numbers of ancestor populations. Furthermore, the results depend on the initialization, also known as *multimodality* (Jakobsson and Rosenberg, 2007; Behr et al., 2016). Challenges such as these increase the risk of over-interpreting the results (Lawson et al., 2018). Several efforts aimed to address specific constraints of ADMIXTURE. For example, Behr et al. (2016) addresses multimodality, while Dominguez Mantes et al. (2023) reimplements the ADMIXTURE framework with neural networks to improve computational efficiency. Moreover, independent of ADMIXTURE, a more recent approach considers a hierarchical structure for four ancestral populations based on the length of ancestral tracts along genomes (Zhang et al., 2024). Methods with a consistent inheritance structure, such as hierarchical clustering methods, may offer benefits in terms of interpreting the inferred ancestries.

Here we introduce tangleGen, a hierarchical clustering method, to infer population structures in population genetics and exploit its capabilities. tangleGen is a model-free approach and does not infer ancestry proportions in the sense of ADMIXTURE and related methods that are based on a model that assumes *K* independent ancestral populations. Instead, it infers a hierarchical clustering of the individuals which sheds light on their ancestral relationships. tangleGen is based on Tangles, a theoretical concept that originated from mathematical graph theory, where they were initially conceptualized to represent highly cohesive structures within graphs (Diestel, 2018, 2019; Diestel and Whittle, 2023).

Recently, the theoretical concept has been translated into an algorithmic framework (Klepper et al., 2023) that offers a highly flexible approach to soft and hard hierarchical clustering. The flexibility of the framework allows for a versatile application for genetic data, where the key components of the algorithm can be customized to account for different data types as well as to focus on different aspects of the underlying population structure. Tangles aggregates information about the data set’s structure from many bipartitions that give local insight. Thereby, the information from many weaker, imperfect bipartitions is combined into an expressive clustering.

tangleGen provides a powerful new tool for researchers in population genetics. It not only improves interpretability by providing a hierarchical perspective on the composition and structure of populations, but also adds a new level of explainability to the clustering by disentangling the individual contributions of relevant SNPs. In addition, tangleGen is deterministic and identifies the number of related populations as a result of its hierarchical clustering, rather than requiring the number of independent populations as a pre-specified input. Thus evolutionary research on species that have a recent and entangled population structure, as well as practical applications where the impact of particular SNPs on the inference is of importance, can specifically benefit from tangleGen.

## Results

The fundamental concept of Tangles revolves around the aggregation of information from bipartitions of the data set. Therefore, tangleGen uses bipartitions of genetic data to achieve a hierarchical soft clustering. We first introduce the concepts of tangleGen using a minimal example before applying the method to both simulated and real data in the next section.

### tangleGen: A Graph-Based Concept for Population Genetics

To infer ancestral relationships, tangleGen proceeds in four steps:

1. *Constructing basic cuts on the set of individuals*: The foundation of tangleGen are many bipartitions on the set of individuals, dividing them into two groups. Thus we will refer to bipartitions as “cuts” here. tangleGen uses cuts based on SNPs, which enhances explainability in the context of population genetics and genetic data.
2. *Assigning costs to cuts to sort them for the hierarchical clustering*: A well chosen cost function favors cuts with higher discriminative power by assigning them lower costs, while penalizing cuts that separate closely related groups of individuals. The correspondingly sorted cuts enable the next iterative hierarchical clustering step.
3. *Iteratively composing the tangles tree by orienting cuts*: Beginning with the lowest-cost cut, the algorithm iteratively orients the cuts to delimit clusters of individuals. Such a meaningful orientation of a subset of cuts that identifies a cluster is called a “tangle”. These tangles form the basis for constructing a tangles tree, which represents the resulting hierarchical cluster structure.
4. *Computing a soft clustering based on characteristic cuts to infer ancestral relationships*: Characteristic cuts are cuts that define the tangles tree and, consequently, the identified cluster structure. Based on them, the soft clustering is computed, a value between 0 and 1 for each individual and cluster, indicating how likely an individual belongs to a particular cluster. Finally, the method infers the ancestry of individuals based on the soft clustering results.

1Let us explain the steps in more detail, based on the following minimal example. We consider nine individuals from three populations. Population “A” comprises four individuals, while populations “B” and “C” comprise two and three individuals, respectively, and are more closely related. We consider six SNPs, and the corresponding genotype matrix is shown in Fig. 1 A.

**Fig. 1.**
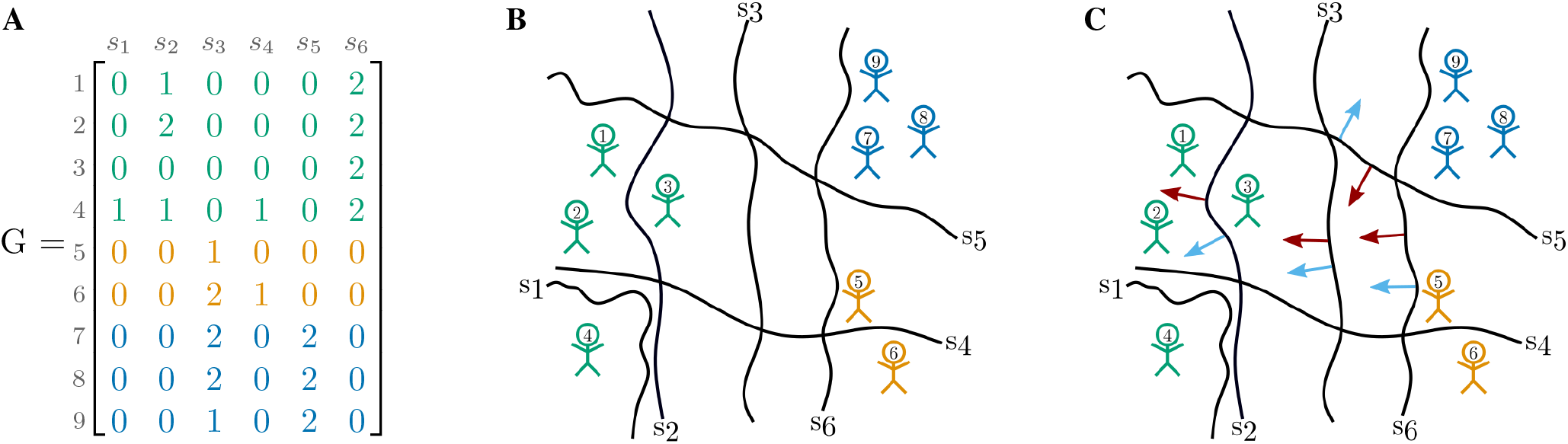
**A:** Representative genotype matrix, which indicates the number of derived alleles per individual per site. Individuals in the rows, SNPs in the columns. **B:** SNPs induce cuts (black lines) on the set of individuals, separating individuals without the derived allele (genotype matrix entry ^0^) from those carrying at least one derived allele (genotype matrix entry *>* 0). Individuals are shown as stick figures with the corresponding individual IDs. The colors correspond to the underlying population structure. **C:** Red arrows indicate a meaningful orientation, that is a consistent orientation of the cuts for agreement parameter *a* = 2, blue arrows a non-meaningful/inconsistent orientation.

### Constructing basic cuts on the set of individuals

The basic ingredients, necessary to construct tangles, are cuts that divide the set of individuals. Naturally, most cuts will not separate distinct populations perfectly from one another. However, the information from many weaker, imperfect cuts is aggregated into an expressive clustering. Therefore, this approach can be seen as a form of boosting (Klepper et al., 2023). For population genetic data, each SNP naturally divides the individuals into two groups: Those individuals who do not carry the derived allele at all (genotype matrix entry is 0) and those who carry it at least once (genotype matrix entry is 1 or 2). A visualization is shown in Fig. 1 B. The rationale behind this procedure is that individuals in the same population are more likely to exhibit similar presence/absence patterns for mutations with discriminative power. Conceptualized in terms of cuts, individuals from the same population tend to lie on the same side of many cuts. The information provided by the cuts will later be aggregated by *tangles*.

### Assigning costs to cuts to sort them for the hierarchical clustering

Clearly for some SNPs the corresponding cuts provide more information about the underlying population structure than others. Even if a SNPs occurs in different frequencies in different populations, the corresponding cut does not necessarily have to distinguish the populations well. But some mutations do distinguish populations, for example mutations in the LCT gene associated with lactose tolerance distinguish very well between Northern Europeans and East Asians (Sahi, 1994; Ingram et al., 2009). To allow tangleGen to exploit this knowledge, we need to provide a cost function to evaluate and sort the cuts. Here, we define a custom cost function to infer demographic structures from genomic data based on the Fixation Index (FST) *F*, a commonly used measure of genetic differentiation. *F* compares the genetic variability within and between populations and is therefore an intuitive choice.

The FST value is usually calculated based on a single SNP and known populations S and T. However, in our case we do not yet know the underlying populations. Instead each cut based on a SNP *s* provides a suggestion on how to separate the individuals into two groups. If the resulting groups *A* and *B* are the underlying populations for the FST calculation, the FST value for the particular SNP *s* is obviously not informative, as it induced *A* and *B*. Therefore, we instead evaluate a cut by taking the mean FST value over all SNPs with respect to the division induced by the cut *s*:

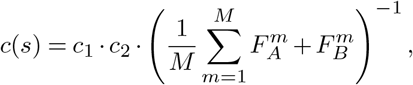

where *M* is the number of SNPs, 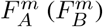the FST value of SNP *s*_*m*_ regarding the separation of group *A* (*B*) and *c*_1_, *c*_2_ are penalizing factors as described in the Methods section, Eq. 3 and Eq. 4. The cost function favors cuts that divide individuals into groups that are more differentiated. The lower the cost, the more expressive the cut for the underlying population structure. However, such a FST-based cost function does not necessarily punish cuts that separate smaller sub-populations if the global division into *A* and *B* still has a high mean fixation index. To disfavor such cuts, which can potentially create inconsistencies early on, we add the penalizing factor *c*_1_ based on k-nearest-neighbors (kNN). Hereby, the number of neighbors to be considered *k* is introduced as a hyperparameter. Supplemental Note 11 includes an analysis of the effect of *k*. The more a cut separates closely related individuals, the more it is penalized by *c*_1_. Second, when inferring population structures using a hierarchical method, it is desirable to identify larger populations first. Therefore, the second penalizing factor *c*_2_ slightly prefers cuts that divide individuals into groups of equal size. Details can be found in the Methods section.

In the minimum example, with *k* = 2 for the kNN penalty *c*_1_, we obtain the following ordering starting with the cut with the lowest cost: *s*_3_, *s*_6_, *s*_5_, *s*_2_, *s*_1_, *s*_4_. The algorithm then processes all cuts sequentially, starting with the most useful cut, *s*_3_.

Low-cost SNPs will be positioned higher in the hierarchical tangles tree and therefore contribute to the formation of the main clusters. Conversely, more costly SNPs are considered later and contribute to more fine-grained population structures or are disregarded in the process.

Of course, the choice of the cost function is not unique. The flexibility of the Tangles framework allows tangleGen to prioritize different aspects by tailoring the cost function. Depending on the specific use case, an alternative cost function to the one presented in this context may be more appropriate. For an alternative cost function based on the mean deviation from a Hardy-Weinberg equilibrium, see Supplemental Note 6.

### Iteratively composing the tangles tree by orienting cuts

Tangles, and therefore tangleGen, aggregates the information provided by the individual cuts by assigning orientations to each set of cuts. Multiple orientations that point towards the same “dense structure”, in our case a group of more closely related individuals, are considered meaningful. The intuition is that orientations can characterize clusters, but not every orientation of cuts characterizes a cluster. For a set of oriented cuts to be meaningful, the intersection of individuals pointed to must contain at least *a* individuals, for each triplet of oriented cuts. Here, *a* represents the agreement parameter, a hyperparameter of tangleGen that roughly defines the minimum size of a cluster to be considered and should be set appropriately for each application scenario. A meaningful orientation of cuts is then called a *tangle*.

Fig. 1 C illustrates this concept for the cuts *s*_2_, *s*_3_, *s*_5_ and *s*_6_ and two possible sets of orientations indicated by red and blue arrows. Let us first consider the set of orientations indicated by the red arrows. This set of orientations is considered meaningful for agreement parameter *a* = 2, and therefore forms a tangle, because for each triplet of orientations (red arrows), all three orientations point to at least 3 *> a* individuals. Conversely, the orientation indicated by the blue arrows is deemed not meaningful: For each triplet of blue arrows containing the oriented cut *s*_5_, the intersection of individuals pointed to by the corresponding blue arrows is empty and thus smaller than *a*. This concept of meaningful orientations outlined here is formalized in the definition of consistency in the Methods section.

Since the number of possible sets of cuts is large and checking the orientations for consistency can become computationally infeasible, tangleGen uses a tree-based search algorithm to handle the complexity. This approach iteratively generates the tangles tree: the cuts sorted by cost are evaluated one after the other, starting with the lowest-cost cut, and each cut is added to the tangle tree if it can be consistently oriented with respect to the previous cuts and their orientations. For an existing tangle, adding an oriented cut can either lead to a larger meaningful set of oriented cuts or to a loss of consistency. For an inconsistent orientation of cuts, consistency can not be restored by adding any cut. Consequently, larger tangles can only be built from smaller tangles, and thus naturally allow for a hierarchical representation in terms of a binary tree (Fig. 2 A).

**Fig. 2.**
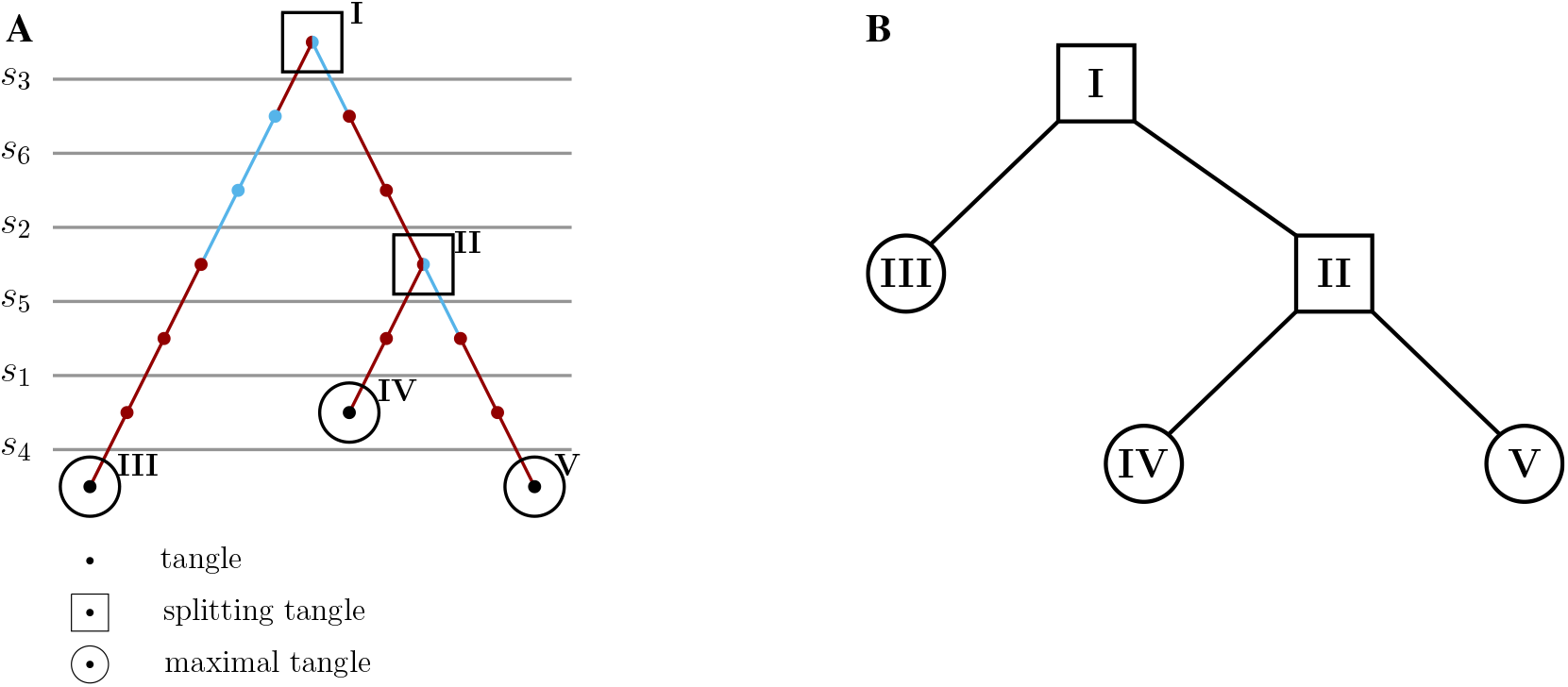
A: Tangles search tree. The constructed tangles search tree for the minimal example for agreement parameter *a* = 2. Cuts, sorted by cost, are evaluated sequentially, starting with the lowest-cost cut (^*s*^_3_). A cut is added to the tangles tree if it can be consistently oriented with respect to the previous cuts and their orientations, forming a tangle. The color indicates whether the orientation of the last added cut points to all individuals with at least one derived allele (light blue) or in the opposite direction (red). When a cut can be oriented in both directions (^*s*^_3_ and ^*s*^_5_), it creates a split at that node, resulting in a splitting tangle (squares). The cut ^*s*^_4_ cannot be added to every branch of the tangle tree. If no cut can be consistently added at any node or all cuts have been added to the tangles tree, the tangles tree is considered complete. The leaves represent the maximal tangles (circles). **B: Condensed tangles tree**. Condensed tangles tree for agreement parameter ^*a* = 2^, that is the tangles search tree from A reduced to splitting and maximal tangles.

Here we illustrate the construction of such a tangles tree along the steps in the minimal example from Fig. 1 As depicted in Fig. 2 A. Starting with the cut of the lowest cost, *s*_3_, the algorithm attempts to meaningfully orient this cut in both directions. For agreement parameter *a* = 2, *s*_3_ can be oriented in both directions, leading to a split in the tree at the root. The next cut with the second lowest cost is *s*_6_. This cut can be oriented with both branches, but only in one direction each, since the orientation must sufficiently overlap with the one of *s*_3_, otherwise their intersection would point to 0 *< a* individuals. Therefore, the existing branches are extended without adding a split in the tangles tree. The same applies to *s*_2_. The cut *s*_5_ can again be meaningfully oriented in both directions, but only if the previous cuts point towards individuals 5, 6, 7, 8 and 9 (as illustrated in Fig.1 C).

Therefore, a split occurs only in the right subtree. As soon as the next cut can no longer be oriented consistently, the algorithm automatically terminates, and the tangles tree is complete. Each node in the tangles tree corresponds to the tangle resulting from the oriented cuts above the node. Consequently, the algorithm alone determines the number of considered cuts and the number of populations. Unlike ADMIXTURE, we do not need to specify the number of populations; we define the size *a* of the smallest group considered a meaningful population. The resulting tree has several levels, where each level 𝓁 represents a step in the hierarchical clustering, and the corresponding nodes give the number of populations identified at that stage. For visualization, see Fig. 3. Additionally, tangleGen offers a pruning option that allows to exclude the external branches in the tangles tree, which are not supported by a sufficient number of cuts/SNPs.

**Fig. 3.**
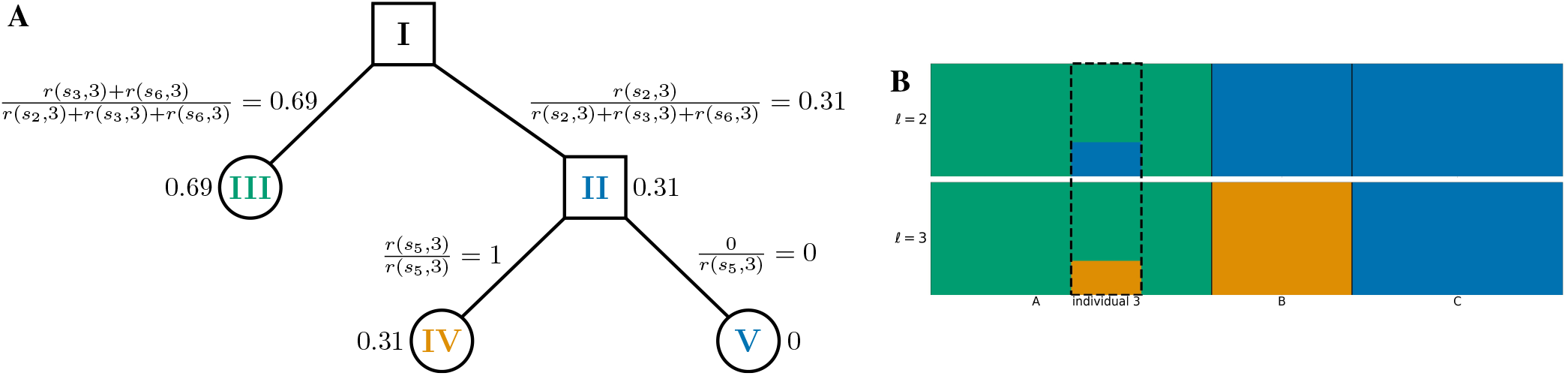
**A:** Soft clustering for individual 3 and agreement parameter *a* = 2. *r*(*s*, 3) refers to the reliability factor for SNP _*s*_ and individual 3 as defined in Eq. 5 in the Methods section. **B:** Hierarchical clustering of the individuals in the minimal example based on the soft clustering, with agreement parameter *a* = 2. Each subplot corresponds to a level 𝓁 in the tangles tree (Fig. 2 B), where the different levels result from splits in the tangles tree. Individuals are shown on the x-axis, the y-axis illustrates the soft clustering per individual. First bar plot shows clustering into two population, second into three. Individual 3 is highlighted because the soft clustering in A is calculated for this individual. tangleGen clusters the individuals hierarchically into the three populations *A, B* and *C*. The first split, which corresponds to the upper bar plot, is supported by three SNPs (*s*_2_, *s*_3_ and *s*_6_) and the second by one SNP (*s*_5_).

In the next step, the constructed tangels tree will be used to cluster the individuals.

### Computing a soft clustering based on characteristic cuts to infer ancestral relationships

Not all SNPs differentiate between different populations — but tangleGen can identify the SNPs that directly contributed to the splits in the tangles tree, which we refer to as characteristic SNPs here. This enhances the explainability of the method. With the FST-based cost function, the characteristic SNPs have a high, but not necessarily the highest, FST value, as the cost function is based on the induced mean FST of all *other* SNPs and takes further aspects into account. For a more detailed analysis, see Supplemental Note 8. In the minimal example, *s*_2_, *s*_3_ and *s*_6_ are characteristic for the first split and *s*_5_ for the second. These characterizing SNPs/cuts are then processed via the soft clustering (Fig. 3 A for individual 3) into the desired inferred ancestry (Fig. 3 B).

The soft clustering step of Tangles can and should be adapted to the application, as it combines all the components of tangles considered so far into the final output, in our case, the inference of ancestral relationships. We thus developed an ancestry-inference-specific soft clustering step. All details can be found in the Methods section. In the following, we explain the concept and intuition behind tangleGen’s soft clustering. This soft clustering is then illustrated in a stacked bar plot. Given the similarity of STRUCTURE and ADMIXTURE plots to this soft cluster bar plot (Fig. 3 B and all further soft clustering plots), it is important to clarify that tangleGen illustrates a hierarchical soft clustering instead of ancestry proportions derived from a probabilistic model with independent populations. In particular, the soft clustering is solely based on characteristic SNPs weighted according to their reliability factor.

To infer the ancestry of the individuals, our objective is to determine, for each individual and split, a value between 0 and 1, indicating how likely, based on the SNP-based cuts, a specific individual belongs to the left or right subtree/cluster. To achieve this, we evaluate each split in the tangles tree and its characterizing SNPs separately for each individual. Given an individual and a split, we could simply compute the fraction of characterizing SNPs that assign the individual to, say, the left subtree/cluster, relative to the total number of characterizing SNPs. However, not every characterizing SNP is equally reliable for identifying population structure. Well-mixed populations are assumed to be in Hardy-Weinberg equilibrium, but SNP-based cuts cannot account for this. For example, the cut based on SNP *s*_2_ in Fig. 1 B separates individuals 1, 2 and 4 from 3, 5, 6, 7, 8 and 9. But, under the Hardy-Weinberg assumption, it is clear that individual 3 is misclassified and should be grouped with individuals 1, 2 and 4 instead. Due to the nature of SNP-based cuts, such misclassifications are inevitable at SNPs that are not homozygous for all individuals, that is the SNPs *s*_1_, *s*_2_, *s*_3_ and *s*_4_ in the minimal example. To alleviate this, we assign a reliability factor to each cut, which calculates the probability that the individuals are correctly assigned to the two sides in respect to an assumed Hardy-Weinberg equilibrium. For explicit calculations, see the section about the soft clustering in Methods. Here we only give an intuition based on cut *s*_2_ in the minimal example.

Considering the side of *s*_2_ with individuals 1, 2, and 4, it is clear that under the assumption of a Hardy-Weinberg equilibrium one homoyzgous sample, here likely individual 3, has been misclassified. Thus, 1 out of 6 individuals on the other side of the cut is expected to be misclassified, resulting in a reliability of 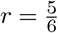For the other side of the cut, we expect at least one of the individuals 1 and 4 to be misclassified, with individual 2 most likely being correctly assigned. This leads to a reliability of about *r* = 0.39. With this, we can now calculate the soft clustering: we take the fraction of characterizing SNPs that assign an individual to one side, and then weight each characteristic cut according to its reliability *r*. Fig. 3 A shows the complete soft clustering of individual 3, which matches our intuition. For example, for the first split, individual 3 has a soft clustering probability of 0.69 for belonging to the left subtree (cluster *A*) and 0.31 for the right subtree (clusters *B* and *C*), which is reasonable as only SNP *s*_2_ orients individual 3 towards the right cluster. Computing the soft clustering for every individual and split results in a hierarchical plot showing the inferred ancestries for each individual, Fig. 3 B. In particular, individual 3 is partially assigned to populations A and B/C, which matches Fig. 1 B. In comparison, due to the very low number of SNPs the minimal example represents a greater challenge for ADMIXTURE, as can be seen in the Supplemental Note 1.

### tangleGen Infers Population Structures

#### Inferring simulated population structures

First, we demonstrate tangleGen on simulated data, which allows the underlying demographics to be specified while controlling for factors such as migration, mutation rate and recombination rate. We used msprime (Baumdicker et al., 2022) to simulate 800 diploid individuals, with a total of 5041 biallelic SNPs within the predefined population structure shown in Fig. 4 A. Details on the simulation can be found in the Methods section *Data Simulation*. Fig. 4 D shows the hierarchical soft clustering of tangleGen with this data set. Clearly, tangleGen clusters individuals into the correct populations. The comparison of Fig. 4 A with Fig. 4 B demonstrates that tangleGen reveals a hierarchical clustering consistent with the underlying population structure in the simulation. However, at the lowest level, involving migration between populations A and E, tangleGen splits A from B before E from F.

For comparison, we run ADMIXTURE with the same simulated data set for *K* between 2 and 12, the input parameter indicating the number of populations to be identified. Results for this simulated scenario, along with those for the 1000 Genomes dataset, using fastStructure and SCOPE, are provided in Supplemental Note 9. In contrast to tangleGen, ADMIXTURE, Fig. 4 C, also correctly assigns individuals for *K* = 8 to their respective populations but, as expected, does not produce a consistent hierarchical clustering. Inconsistencies are, for example, already observed between *K* = 2 and *K* = 3, where populations ABCD and EFGH are initially determined as ancestral populations, and in the next level, ABH, CD, and EFG, leading to an arrangement inconsistent with any underlying demography. Furthermore, interpretability with ADMIXTURE is hindered by the uncertainty surrounding the choice of the value of *K*, an input parameter. This makes it unclear which *K* value best captures the most detailed population structure. From *K* = 9 ADMIXTURE does not determine any further populations, but also already subdivides population H at *K* = 6. In contrast, tangleGen demonstrates consistency, stopping when it cannot identify additional populations of sufficient size, and is unaffected by random factors, unlike ADMIXTURE, which exhibits multimodality issues as the inferred ancestry proportions vary significantly depending on the initialization.

**Fig. 4.**
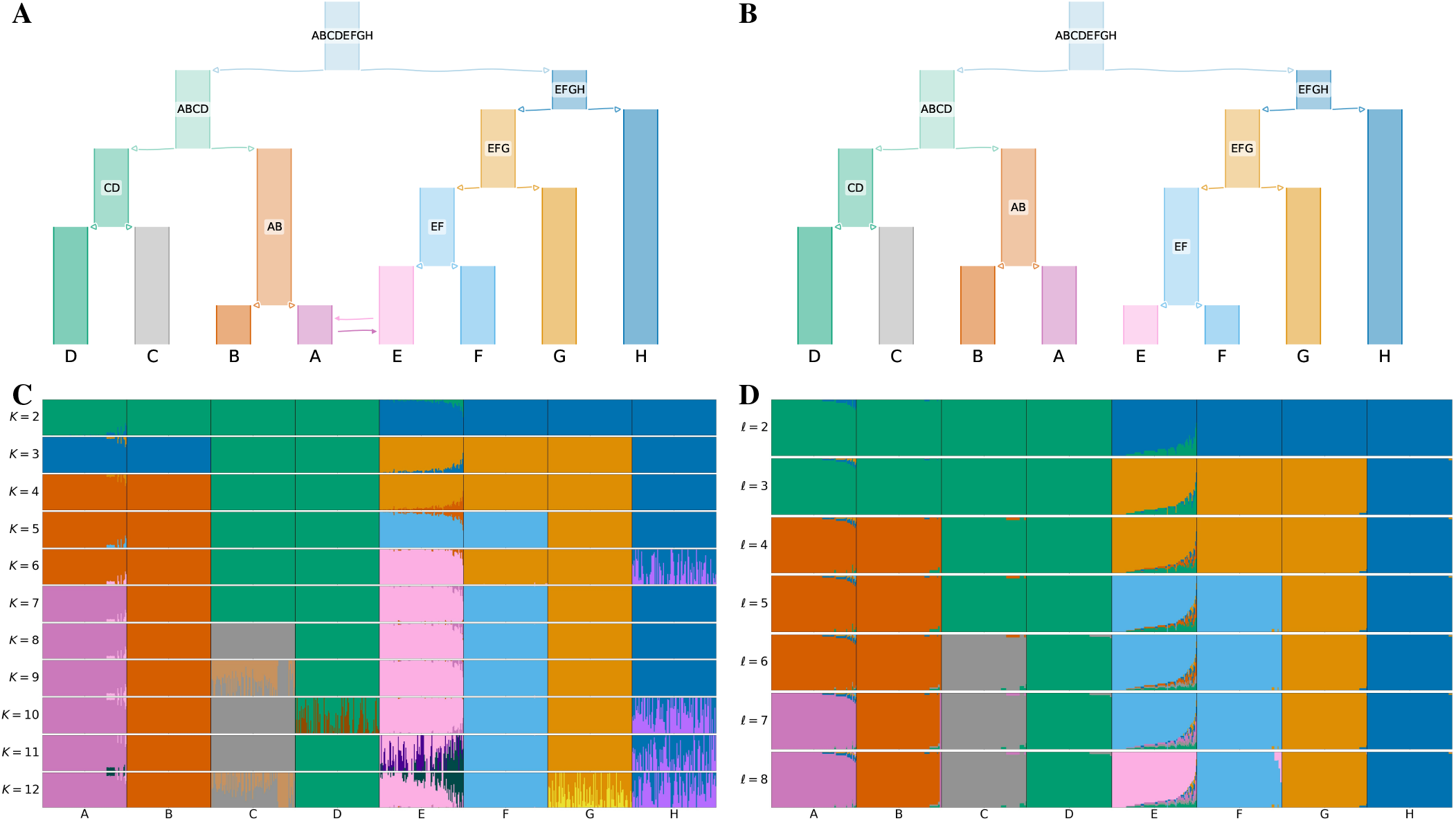
**A: Underyling demographic structure in simulation** with migration added between populations A and E. **B: Inferred population structure by tangleGen**. The hierarchy extracted from the tangles tree in tangleGen coincides with the simulated population structure except for the reversed order of splitting AB and EF. Note that tangleGen only estimates the population structure and not the height of the hierarchy; the branch lengths shown here are purely schematic. Figures A and B are visualized with *DemesDraw*. **C: ADMIXTURE ancestry proportions**. Inferred ancestries by ADMIXTURE from simulated genetic data for different numbers of populations, denoted by *K*. Individuals are sorted within the populations in the same order as in the tangleGen soft custer plot D. The best of ten runs is shown in terms of clarity of clusters and minimization of inconsistencies compared to the underlying population structure. ADMIXTURE estimates the ancestry proportions well for the true number of populations *K* = 8, but gives also plausible results for other choices of *K*, although for *K >* 8 the simulated data does not contain any further population structure. Furthermore, the results for different *K* show inconsistencies, for example, for *K* = 3 and also *K* = 6. **D: Soft clustering of tangleGen**. Inferred ancestral relationships by tangleGen on simulated data with underlying demography as shown in A. The individuals are sorted within the populations based on the soft cluster proportions to achieve a block structure in the plots. Each subplot corresponds to a level 𝓁 in the tangles tree, where the different levels result from splits in the tangles tree. tangleGen clusters individuals hierarchically into the correct populations, whereby the hierarchy matches the underlying population structure up to the lowest level. In contrast to ADMIXTURE, tangleGen independently determines the depth of the population structure and completes the inference when no further SNP-based cuts can be consistently added. Starting with the top split between the ancestral populations ABCD and EFGH, each population split is based on 51, 177, 96, 76, 139, 1 and 18 characteristic SNPs. tangleGen parameters: Agreement parameter *a* = 50 and cost function as in Eq. 2 with *k* = 40.

tangleGen is particularly effective at inferring ancestral relationships from clearly differentiated populations, as demonstrated by these simulations. However, tangleGen also handles more complex demographies well, such as those with increased migration leading to less differentiated populations. Fig. 5 illustrates tangleGen’s performance on simulations with a quadrupled migration rate and migration between all populations from A to H. Due to tangleGen’s hierarchical structure, the migration patterns are clearly recognizable. In addition, except for an early split of population E, which shares ancestry with both subtrees in the underlying population structure (Fig. 5 A), the order of splits is consistent with the underlying population structure. Population E is also already split off using a low *K* with ADMIXTURE (see Supplemental Fig. S10 D). Furthermore, increased migration reduces the number of SNPs that separate populations well. This is also reflected in tangleGen, as clustering is now based on fewer SNPs. If the migration rate is increased even further, this effect becomes more pronounced, and the barplot of the soft clustering becomes less smooth (see Supplemental Fig. S10). A detailed analysis with additional migration scenarios, including a comparison to ADMIXTURE, is provided in Supplemental Note 10. Applications that involve a much higher number of SNPs pose a greater challenge to tangleGen, as expected for any hierarchical SNP-based method. For an example involving significantly more SNPs and significantly shorter time intervals between population splits, see Supplemental Note 6.

**Fig. 5.**
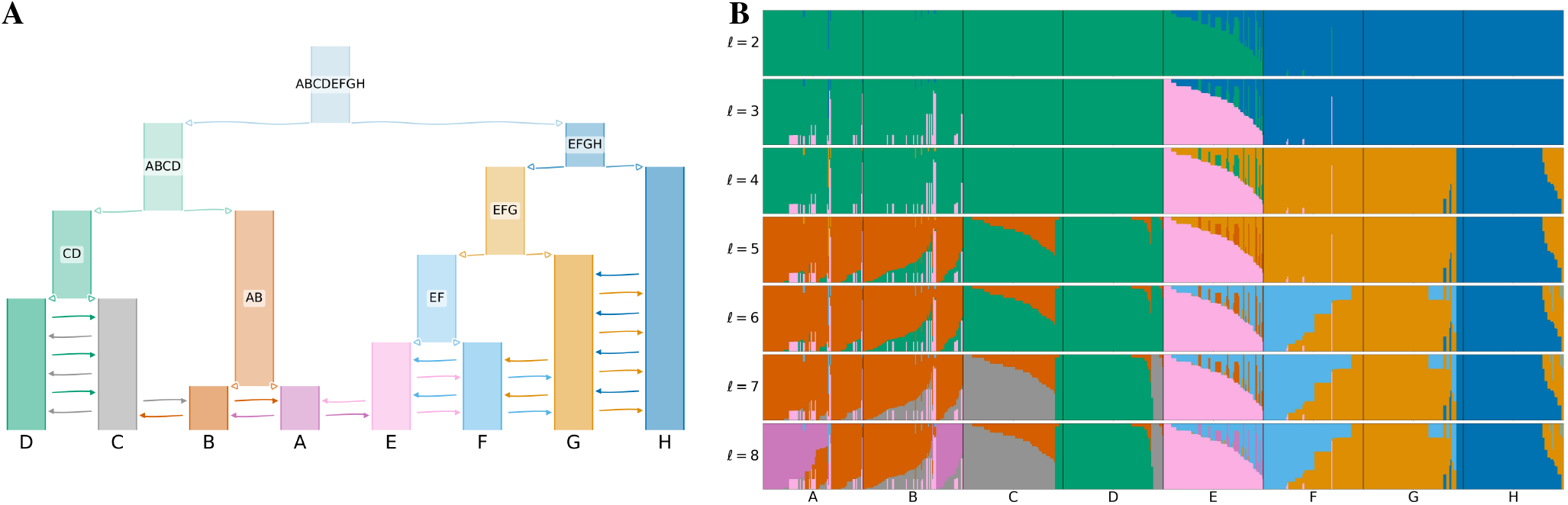
**A: Underlying demographic structure in the simulation** with migration added between populations A to H as indicated by the arrows. Migration rate is quadrupled in comparison to Fig. 4 A. See the Data Simulation section in Methods for more details. **B: Soft clustering of tangleGen**. Inferred ancestral relationships by tangleGen on simulated data with underlying demography as shown in A. The plot shows the soft clustering for different levels 𝓁 just as in Fig. 4 D. Note, that the maximal value of 𝓁 is a result of its hierarchical clustering in combination with the agreement parameter. tangleGen infers meaningful population structures even with increased migration rate. Higher migration rates result in more admixed populations and the hierarchical nature of tangleGen shows the migration pattern clearly. Starting with the top split between the ancestral populations ABCDE and FGH, each split is based on 29, 11, 10, 36, 10, 4 and 3 characteristic SNPs. In contrast to Fig. 4 D, the agreement parameter is set to *a* = 30 and all external branches supported by only one SNP are pruned (pruning parameter equals 1), to account for the less differentiated populations.

#### Inferring population structure within the 1000 Genomes Project

We used 2157 diploid individuals sampled in the data from 1000 Genomes Project Consortium et al. (2015) Phase 3 and extracted the SNPs of the AIMs panel by Kidd et al. (2014). This panel is widespread, includes 55 SNPs strategically distributed across almost all chromosomes, and is known to be suitable for distinguishing superpopulations within the 1000 Genome data set: Africa (AFR), Europe (EUR), South Asia (SAS) and East Asia (EAS) (Pfaffelhuber et al., 2020).

Fig. 6 B shows tangleGen’s soft clustering for Kidd’s AIMs panel. It is evident that tangleGen correctly clusters the individuals into the four superpopulations: African (AFR), European (EUR), South Asian (SAS), and East Asian (EAS) populations. The inferred hierarchy reflects meaningful patterns and aligns with the out of africa dispersal (Mellars, 2006). Fig. 6 A illustrates that individuals that are not assigned to a single population by tangleGen, are the ones positioned between multiple populations within a PCA analysis based on the AIMs panel. Furthermore, once again, the hierarchical nature of soft clustering is evident, and the leaves of the tree, when tangleGen has inferred the maximal level 𝓁 = 4, can be assigned to the AFR, EUR, SAS, and EAS superpopulations from left to right. This soft clustering is then processed into the plot in Fig. 6 B.

**Fig. 6.**
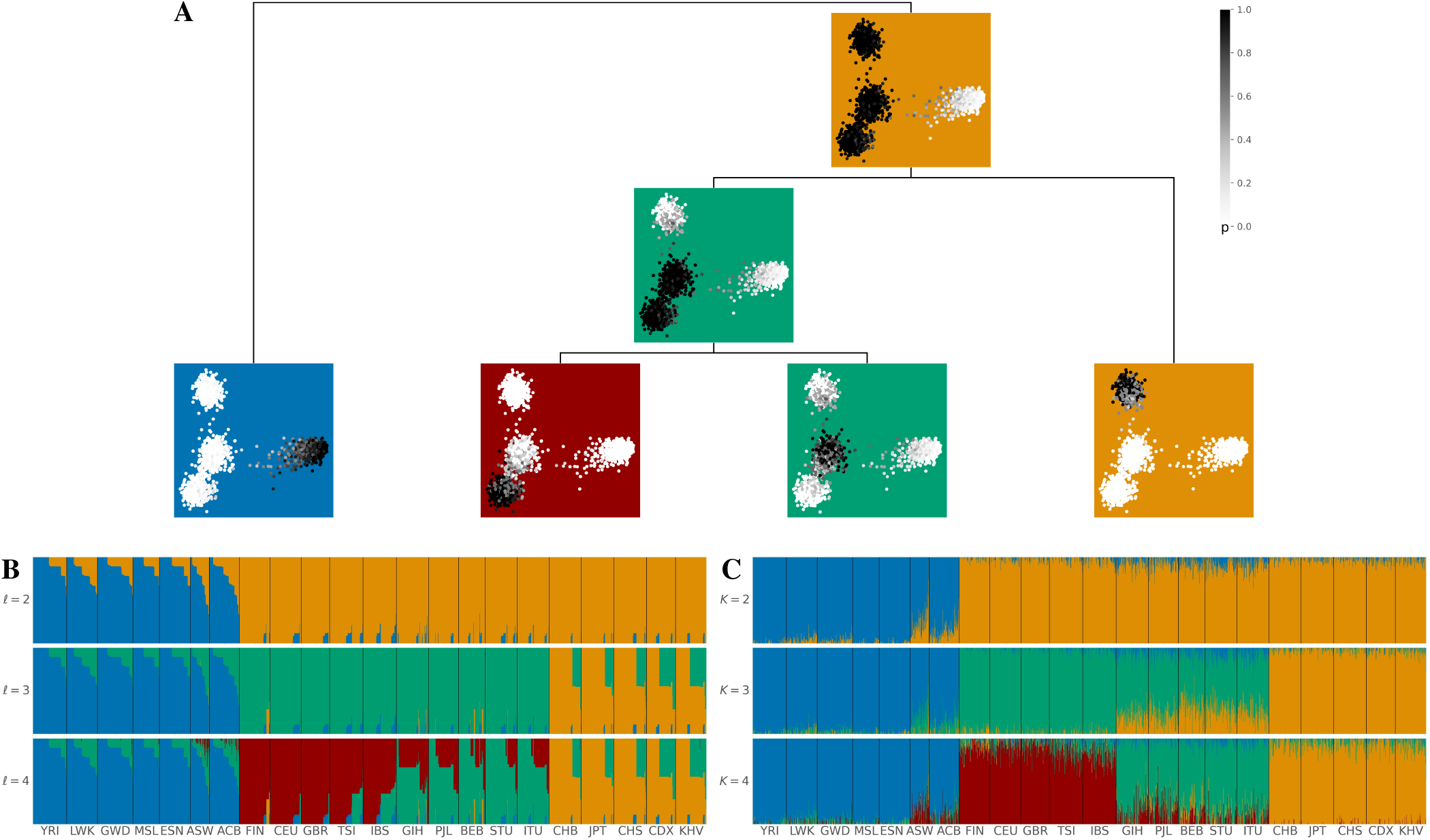
Inferred ancestries by tangleGen and ADMIXTURE based on SNPs from Kidd’s AIMs set for individuals sampled in the 1000 Genomes Project Phase 3, excluding AMR. **A: Soft clustering of tangleGen visualized with PCA** for each individual and each split in the tangles tree. Panels at each split in the tangles tree show a PCA plot based on the AIMs set genotype matrix. Each individual is represented as a dot, with darker colors indicating higher confidence in the assignment of the individual to the corresponding cluster. The PCA visualization highlights that individuals positioned between two populations are softly clustered into both populations. The same soft clustering is processed in B into an ADMIXTURE like bar plot. The resulting soft clustering is hierarchical and the leaves of the tree can be assigned to the AFR (blue), EUR (red), SAS (green), and EAS (orange) superpopulations. **B: Soft clustering by tangleGen visualized in a bar plot**. Each subplot corresponds to a level 𝓁 in the tangles tree, where the different levels result from splits in the tangles tree. tangleGen identifies the four superpopulations AFR, EUR, SAS and EAS with a meaningful inferred hierarchical population structure. The first split is supported by nine characteristic SNPs, and the second and third by three characteristic SNPs each. Cost function as specified in Eq. 2 with *k* = 40, and agreement parameter *a* = 225. **C: Ancestry proportions by ADMIXTURE**. ADMIXTURE identifies the four superpopulations AFR, EUR, SAS and EAS, when the number of populations is set to *K* = 4, but shows inconsistencies, compared to smaller values of *K* in the differentiation between the four populations. The best of ten runs is shown in terms of the clarity of the inferred populations and minimization of inconsistencies compared to the underlying population structure. In B and C, the individuals are sorted within the populations in the same way such that a block structure for the tangleGen plot B is achieved. The four superpopulations are made up as follows: AFR: YRI, LWK, GWD, MSL, ESN, ASW, ACB; EUR: FIN, CEU, GBR, TSI, IBS; SAS: GIH, PJL, BEB, STU, ITU; EAS: CHB, JPT, CHS, CDX, KHV. Results that include Ad-Mixed American (AMR) populations can be found in the Supplemental Note 2.

For comparison, we run ADMIXTURE with a fixed number of ancestral populations between 2 to 4, as shown in Fig. 6 B. Like tangleGen, ADMIXTURE also identifies the four superpopulations, with tangleGen clustering in a similar way to the well-established method. However, as in the simulation, ADMIXTURE also shows inconsistencies here, even if these do not relate to miss-classifying entire populations. For example, when differentiating into three populations, we observe that the South Asian populations share a significant part of their ancestry with the East Asian populations. However, when differentiating into four populations, this component disappears almost completely. Again, ADMIXTURE’s performance depends on the choice of *K*. Fig. 6 considers *K* only up to 4, which is reasonable in this application as ADMIXTURE starts to identify non-existing population structures from there (see Supplemental Note 3 for *K* up to 7). In contrast, tangleGen determines the number of populations as a result of its hierarchical clustering in combination with the agreement parameter.

#### Identification of characteristic SNPs in the tangles tree

Apart from the inherent advantages of a hierarchical method, tangleGen offers the possibility to identify specific cuts and therefore SNPs crucial for the cluster formation in the tangles tree. This enhances the method’s explainability, as identified SNPs can be cross-referenced with expert knowledge. tangleGen determined 15 out of 55 SNPs of the Kidd’s AIMs set as characterizing SNPs to cluster the 2157 individuals in the 1000 Genomes Project (AMR excluded), Fig. 6 A. In particular, nine SNPs were found to be characteristic for the first split in the tangles tree separating the AFR populations, while three each were used for the second and third splits that separate the EUR, SAS, and EAS populations. The characterizing cut with the lowest cost, SNP with ID rs2814778 (following dbSNP ID notation by Sherry et al. (2001)), influences all subsequent orientations and therefore determines that AFR populations are identified first in the hierarchical tangles tree. Notably, this SNP is a recognized component of the molecular basis of the Duffy blood group system, linked to malaria resistance and exhibits near-fixed differentiation between Africans and the other superpopulations (Pfaffelhuber et al., 2020). Furthermore, rs3827760 is known to separate the East Asian populations from the other superpopulations (Pfaffelhuber et al., 2022) and has been identified by tangleGen as the main SNP responsible for the corresponding split in the tangles tree. Finally, rs1426654, a SNP associated with iris color and skin pigmentation in South Asians (Riddell et al., 2020), is the lowest-cost characteristic SNP for the third split in the tangles tree separating between European and South Asian populations. Although tangleGen aims to extract the genetic background structure and does not explicitly identify or require SNPs under local adaptation, such SNPs will accelerate the clustering process and thus frequently appear among the most reliable characterizing cuts. In total, all 15 SNPs identified by tangleGen are meaningfully linked to their respective population splits in the tangles tree. The complete list of characterizing SNPs is provided in Supplemental Note 4.

In addition, tangleGen can be employed to construct a cluster-typical set of SNPs, comprising relevant cuts for each cluster alongside their orientations. This can be useful as it, for example, facilitates a rapid, albeit approximate, clustering of individuals without the need to rerun the method. Note, the cluster-typical SNPs do not represent the genome of any specific individual as consistencies during the construction of the tangles tree are always only ensured for triplets of cuts. Instead they represent the abstract concept of a cluster center. Supplemental Note 5 includes the cluster-typical SNPs for the application on the 1000 Genomes Project data based on Kidd’s AIMs panel.

## Discussion

### tangleGen leverages the hierarchical Tangles framework for population genetics

Here we present tangleGen, an ancestry inference method that exploits the flexibility and robust hierarchical functionality of the Tangles framework. Robustness comes from the fact that Tangles provides results that are deterministic and consistent across different numbers of populations and do not exhibit multimodalities. The flexibility of Tangles lies mainly in the customization of the cuts and cost function to the specific research questions and data structures. This allows tangleGen to focus on ancestry inference based on sequencing data. tangleGen investigates the ancestry of diploid individuals based on biallelic SNPs using cuts that separate individuals according to the presence or absence of at least one derived allele. This one-to-one relationship between SNPs and cuts provides interpretability, as it allows identifying individual SNPs responsible for the inferred population structure. However, as the derived allele is usually not homozygous for all individuals in any subpopulation, the corresponding cut will inevitably misassign some individuals. This error is mitigated by the weights of the cuts in the soft clustering. To obtain a meaningful hierarchical structure within the tangles tree we need to prioritize cuts according to their discriminative power between populations. Therefore, it is crucial to assign a suitable cost to each cut. Here, we opt for a cost based on the fixation index, a classical approach to distinguish populations. To derive meaningful inferred ancestral relationships from the tangles tree, tangleGen accounts for the reliability of the cuts. To do so, the soft clustering incorporates an estimate of the number of misclassified individuals according to an assumed Hardy-Weinberg equilibrium.

The Tangles framework has the flexibility to emphasize different research questions as incorporating alternative definitions of cuts, cost functions, and soft clustering allows to focus on different aspects of the data. Future investigations could, for example, explore cuts that consider multiallelic sites or separate haploids. Furthermore, cuts based on multiple SNPs rather than single SNPs could improve the performance but might lower the interpretability. Given the fundamental principles underlying FST and Hardy-Weinberg equilibrium (HWE), we expect that the proposed FST-based cost function and HWE-based reliability factor will be suitable for most diploid species. Nonetheless, the costs of the cuts and the soft clustering can be adjusted to specific use cases. As an example we propose an alternative cost function based on the divergence from an Hardy-Weinberg equilibrium in Supplemental Note 6. Together, these features of the Tangles framework make tangleGen a particularly flexible approach in population genetics.

### tangleGen parameters in practice

Through the choice of hyperparameters tangleGen provides an additional option to focus on different aspects of the clustering (Klepper et al., 2023). The agreement parameter *a* roughly corresponds to the minimum size for a cluster/population to be identified. Adjusting the agreement parameter provides the flexibility to obtain either major clusters (with a large *a* value) or minor clusters (with a small *a* value), allowing users to experiment with different settings. Typically, the agreement parameter should be set between 33% and 66% of the smallest desired population (Klepper et al., 2023).

For example, in the simulated scenario, the agreement parameter is set to 50, which corresponds to half of each population size. For the analysis of the 1000 Genomes Project data set in Results, the agreement parameter is set to 225, which corresponds to about 34% of the largest superpopulation (AFR) and 46% of the smallest superpopulation (SAS). Further details are added in the Methods section and Supplemental Note 11.

As Tangles does not require many SNPs for clustering and aggregates the available information efficently, we concentrated on data sets that contain a comparably low number of SNPs, which are the most promising application of the current method. For datasets with a higher number of SNPs, tangleGen currently exhibits reduced speed and accuracy. However, future improvements, such as the implementation of a more efficient cost function, can enhance its performance. Clearly, every hierarchical SNP based method will face unique challenges when dealing with large numbers of SNPs. A too early consideration of a SNP positioned high in the tangles tree may introduce cuts that are not representative of the actual population structures. When dealing with such extensive datasets containing a considerably higher number of SNPs compared to the two scenarios considered here, using the pruning option that defines the minimum number of cuts required to support a split in the tangles tree can help to avoid noise clusters (Klepper et al., 2023). In our experience, a pruning parameter of 1 or 2 is often sufficient. For population structures with less differentiated populations, it is advisable to use a smaller agreement parameter and positive pruning to ensure the detection of the populations while mitigating the formation of noise clusters. Further details can be found in the Supplemental Note 6 and 10.

### tangleGen infers interpretable population structures

To evaluate tangleGen for inferring ancestral relationships, we applied the method to both simulated and the 1000 Genomes Project data. tangleGen performs comparably well to the established ADMIXTURE method in terms of clustering and ancestry inference. The advantage of using tangleGen lies in its interpretability. The hierarchical nature provides insights into ancestral relationships and offers a new perspective on the composition and evolution of populations. Unlike ADMIXTURE, tangleGen operates independently of any random factor and the specified number of ancestral populations *K*. Instead, tangleGen determines the number of populations as a result of its hierarchical clustering in combination with the agreement parameter as the algorithm terminates automatically if the consistency condition is no longer met. At this point, no further populations are split, and the number of maximal tangles in the tangles tree determines the final number of populations. Furthermore, as discussed in the Results section, tangleGen yields the characteristic SNPs for clustering. This increases the explainability of the method and allows to identify SNPs that differentiate between populations in the tangles tree. Concentrating on a smaller set of automatically inferred cluster-distinguishing SNPs can be beneficial for the analysis of large-scale data sets, such as biobanks. In addition, a cluster-typical set of oriented characterizing SNPs can be used to cluster new individuals without running the method again. Identifying an explicit genetic state for each of the populations can provide a more intuitive view of the inferred ancestry. However, it is important to note that this set of oriented SNPs simply represents the center of the identified clusters and does not correspond to any existing individual in the population.

## Conclusion

The hierarchical approach in tangleGen not only infers ancestries and identifies populations, but also constructs consistent relationships between populations over time. In addition to the inherent hierarchy, the ability to define a cluster-typical set of characteristic SNPs responsible for the clustering decisions distinguishes it from other methods. Moreover, tangleGen provides deterministic results and automatically determines the number of populations without requiring the user to specify. Furthermore, tangleGen inherits the flexibility of the underlying method, Tangles. With the ability to customize cuts, cost function, soft clustering and hyperparameters, tangleGen can be tailored to a wide range of research questions and is a versatile tool to investigate the genetic diversity and ancestral relationships between populations.

## Methods

### tangleGen Implements the Tangles Concept for Population Genetics

Tangles originate in mathematical graph theory, where they capture the concept of a highly cohesive structure in graphs. In their original form, they are based on all possible bipartitions of the nodes in a graph, and the existence of a tangle defines a densely connected structure. The algorithm presented in Klepper et al. (2023) distills this concept to its essence to cluster arbitrary data hierarchically. In the following, we outline the key concepts of Tangles and describe the specific adjustments to utilize the framework in the context of ancestry inference, specifically constructing SNP-based cuts, the cost function, and soft clustering. The implementation of tangleGen is available on Github: k-burger/tangles_in_pop_gen.

We use the notation and definitions introduced in Klepper et al. (2023). Assume we are given a set *V* = *{v*_1_ …, *v*_*N*_ *}* of individuals. A subset *A ⊂ V* induces a **bipartition** or **cut** of the data into the set and its complement *S* = *{A, A*^*∁*^*}*. In order to construct tangles, we will consider a set of initial cuts *S* = *{{A*_1_, *A*^*∁*^ *}*, …, *{A*_*M*_, *A*^*∁*^ *}}*.

### Constructing SNP-based cuts to infer ancestries

Since we want to build upon the Tangles framework to analyze ancestries and population admixtures, we follow the most common methods and use the genotype matrix *G* = (*g*_*nm*_) *∈ {*0, 1, 2*}*^*N×M*^ as input, with *g*_*nm*_ the observed number of copies of the derived allele at site *m* of individual *n*. In particular, we consider diploid individuals and biallelic SNPs. To infer ancestries from the genotype matrix, we choose an initial set of cuts induced by SNPs. Thereby, each cut separates all individuals into two sets of individuals based on a single SNP, so that individuals carrying the derived allele at position *m* (*g*_*nm*_ *∈ {*1, 2*}*) are separated from those not carrying the derived allele at position *m* (*g*_*nm*_ = 0).

### Consistent orientations of cuts and tangles

For a single cut *S* = *{A, A*^*∁*^*}*, an orientation “points” towards one of the sides. In our minimal example, SNP *s*_6_ points towards *A* = *{*1, 2, 3, 4*}*, that is, the individuals carrying the derived allele at SNP 6. For a set *S* of cuts, we define an orientation *O*_*S*_ by choosing one side for each cut, giving an orientation to every *S ∈ S*. We write *A ∈ O*_*S*_ if *{A, A*^*∁*^*} ∈ S* and *O*_*S*_ orients it towards *A*. The intuition is that orientations can characterize clusters, but not every orientation characterizes a cluster: the orientations need to be “consistent” in some way. For a meaningful orientation, we have to ensure that the chosen sides of all the cuts point to one meaningful subgroup. This is precisely the purpose of tangles. *Definition 1* (**Consistency and tangles)** Let *S* be a set of cuts on a set *V*. For a fixed parameter *a ∈* ℕ, an orientation *O*_*S*_ of *S* is *consistent* if all sets of three oriented cuts have at least *a* individuals in common:

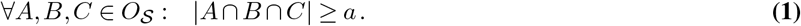

We call Eq. (1) the *consistency condition* and *a* the *agreement parameter*. A consistent orientation of *S* is called a tangle. In the minimal example, Fig. 1 C, we illustrate the consistency condition for the following set of bipartitions:

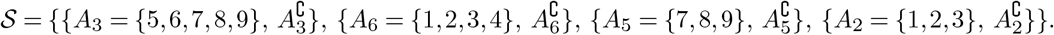

The **agreement parameter** *a* controls the minimum degree to which the orientations have to agree. When *a* is chosen too small, it weakens the induced consistency condition, causing the algorithm to identify non-cohesive subgroups. On the other hand, we want to avoid a too-large *a*; in practice, it should be slightly smaller than the expected size of the smallest cluster to account for noise. If the cuts in *S* respect the cluster structure, and if we have a diverse and rich set of *S*, we can reduce *a* without mistakenly marking disconnected structures as clusters (compare with the appendix of Klepper et al. (2023)).

***Finding all the tangles***

To identify all *tangles* within a set of cuts, the algorithm in Klepper et al. (2023) progressively constructs a tangle search tree by considering cuts in an iterative manner. Useful cuts are prioritized, that is, those that lead to a meaningful split of the individuals (for example, do not cut through a group of similar individuals). This intuition is formalized through a **cost function** *c* : *S →* ℝ. This function, application-dependent, quantifies the “quality” of a cut. Our choice of cost function is discussed in detail in the following section. The resulting labeled binary tree (for example Fig. 2 for the minimal example) contains all consistent orientations (tangles) within the considered subset of cuts that do not exceed some cost. Each node corresponds to a specific orientation of a particular cut, ensuring a one-to-one relationship with tangles. Specifically, for a node *e* with *i* nodes on the root-to-*e* path, the corresponding node labels form a consistent orientation of *S*_1_, …, *S*_*i*_, that is a tangle.

### New cost function for inferring ancestries

Tangles offers the flexibility to tailor the method to various applications and its requirements by using a suitable cost function. In order to infer ancestral relationships with tangleGen, we have chosen a cost function based on the fixation index FST, which has already been introduced in the minimal example in Results and is described in detail below. Let us consider a cut *S*_*m*_ that separates the individuals into groups *A* and *B* = *A*^*∁*^, *T* = *A ∪ B*, with *N*_*A*_, *N*_*B*_ and *N* the group sizes. Since each SNP introduces a cut, we average the FST value *F* over all SNPs rather than considering only the SNP responsible for the cut. Let 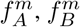 and 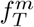 be the frequency of the derived allele at SNP *S*_*m*_ in the groups *A, B* and total population *T* respectively:

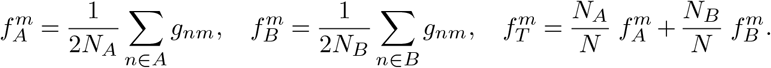

Then, the mean FST value *F* for subpopulation *A* is given by

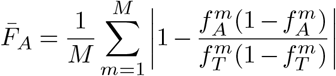

and the resulting cost function *c* : *S →* ℝ is defined by

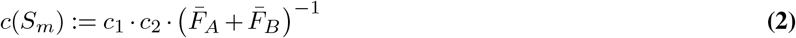

with penalization factors *c*_1_ (Eq. 3) and *c*_2_ (Eq. 4). If kNN_A_in_B is the number of nearest neighbours from A that lie in B and kNN_B_in_A respectively, the penalization factor *c*_1_ is given by

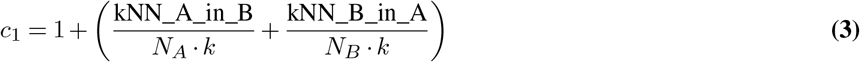

with *k* the number of neighbors considered and

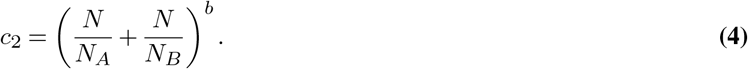

Hereby, *b* weights the penalization factor *c*_2_. The higher *b*, the more severe cuts are penalized that divide individuals into groups of unequal size. In practice, *b* = 0.05 has proven to be a good choice, as cuts that divide individuals into groups of unequal size are sufficiently penalized, but changes in 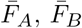and *c*_1_ are still reflected in the cost. The lower the cost *c*(*S*_*m*_), the more informative the cut is for the underlying population structure. To get a better impression of the influence of the cost function, we analyzed the impact of using a random instead of a cost-based order for the cuts (Supplemental Note 12), the effect of the penalizing factor *c*_2_ (Supplemental Note 13), and the hyperparameters *k* and *b* (Supplemental Note 11).

### Computing the clustering

The tangle search tree is constructed hierarchically on cuts, acting as a proxy for our ultimate interest: a hierarchical cluster structure of our individuals. To simplify, we can convert this tree into a condensed form, denoted as *T* ^*∗*^, resembling a dendrogram (shown in Fig. 2 B for the minimal example). Internal nodes, which we call splitting tangles, represent divisions between regions in the data. However, for a single individual, a splitting tangle does not enforce a binary decision; rather, the condensed tree allows assigning probabilities for each individual and tangle, yielding a soft clustering. In the following, we explain how to summarize the information into a condensed tree and how to derive the soft clustering from it.

We first condense the tree to the splitting tangles and only keep the information of cuts that do give insight into the cluster structure; for every split in the tree, we identify all cuts that contribute to the split and thus ‘characterize’ the separation of the two denser structures. In our minimal example, Fig. 2, we consider the cuts *s*_2_, *s*_3_ and *s*_6_ as characterizing cuts for the first splitting tangle *τ* at the node *I* since they provide information about the separation between the left and the right group. For the second splitting tangle *τ* at node *II* we identify *s*_5_, which separates the upper from the lower cluster structure on the right side. All information is taken from the tangles search tree. For a cut *S* to be characterizing for a split, every tangle corresponding to a leaf in the one subtree needs to orient *S* the one way, and every tangle corresponding to a leaf in the other subtree needs to orient *S* the other way. In this sense, the characterizing cuts are the ones that help in distinguishing between the two subtrees of *τ*. In our minimal example, we therefore have a set of characterizing cuts or rather characterizing SNPs *S*(*τ* |_*I*_) = *{s*_2_, *s*_3_, *s*_6_*}* for the first split *I* and for the second *S*(*τ* |_*II*_) = *{s*_5_*}*.

More formally, let *S*(*τ*) be the orientation of *S* in a tangle *τ* and let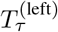 be the left subtree and 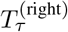 be the right subtree of the node at a tangle *τ*. Then we define the set of characterizing cuts as

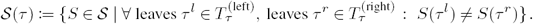

Based on this information, the tree is condensed, and the set of characterizing cuts is tracked for each of the splitting tangles.

### Inferring ancestries from the soft clustering

Let us now compute the soft clustering. Using the characterizing cuts, we derive a heuristic to assess how likely an individual *v ∈ V* belongs to the left or right subtree (or in our case rather a subgroup) of *τ*. For each individual *v* and splitting tangle *τ*, we could simply calculate the fraction of characterizing cuts oriented toward *v* relative to the total number of characterizing cuts. However, not every characterizing cut is equally reliable in separating the identified populations. While characterizing cuts are well suited to distinguish structures based on the mean FST-value, each cut inevitably misclassifies some individuals. Once an SNP is not homozygous for each subpopulation individual, the corresponding cut will make classification errors. To solve this problem, we assign a reliability factor to each cut, which calculates how likely individuals are correctly assigned to the two sides based on the estimated allele frequencies assuming a well-mixed population in Hardy-Weinberg equilibrium on both sides of the cut. For intuition, see the soft clustering section for the minimal example within Results.

For a given cut *S*_*m*_, let 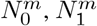 and 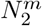 be the number of individuals that carry the derived allele not at all, once or twice. As before, let *A* be the group of individuals that carry the derived allele at least once and *B* = *A*^*∁*^. Then, a first rough estimate of the frequency of the derived allele in *A* is given by

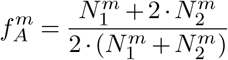

and in *B*, based on the heterozygotes

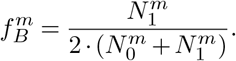

Hereby, 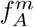tends to overestimate and 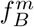tends to underestimate the true allele frequencies. To address this issue, we propose an iterative procedure to approximate the true allele frequencies. Initially, we start with the allele frequencies 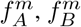. Remember that group *A* consists solely of individuals that have at least one derived allele and all individuals in group *B* are homozygous non-derived. Based on 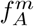and 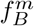we calculate the expected number of individuals that do not carry derived allele and should belong to group *A* and the expected number of individuals that are homozygous or heterozygous for the derived allele and should belong to group *B*:

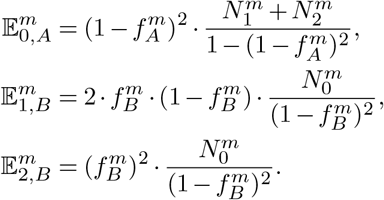

For the next iteration, we then update the estimated allele frequencies

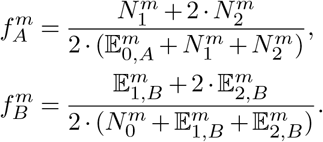

This procedure is repeated until the estimated allele frequencies converge, which is usually achieved after 20 *−* 50 iterations. The final reliability factor of cut *S* and individual *v* is then based on the expected number of misclassified individuals according to the estimated allele frequencies 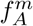 and 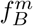:

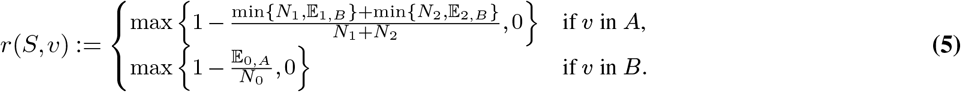

With this, we can now calculate the soft clustering. Let the set *{S ∈ S*(*τ*) | *v ∈ S*^(right)^*}* represent characterizing cuts oriented toward individual *v* on the right side of the tree. Then, we assign a probability *p*^(right)^ for belonging to the right subtree at a tangle *τ* to every individual *v*:

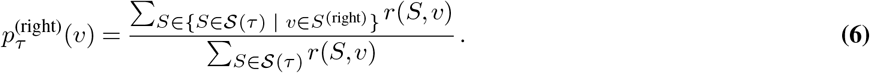

Based on these probabilities, we define the probability *p*_*k*_(*v*) that individual *v* arrives at node *e* as the product of the edge probabilities along the unique path from the root to *τ*. The soft clustering becomes clear by the minimal example and is explained in detail for this example in Results, see also Fig. 3 A. Note that the calculations do not account for a possible linkage disequilibrium (LD) between the SNPs. Pre-filtering SNPs based on LD, as in the Kidds AIMs panel, may therefore be beneficial. However, as SNPs in LD are expected to result in similar cuts, taking them into account could have a small effect on the soft clustering proportions, but does not affect the tangles tree and the resulting inferred hierarchy.

### Computing the hard clustering

If desired, we can now assign individuals to a hard clustering: We assign each individual to the tangle with the highest probability based on the soft clustering.

### Run time of tangleGen

In terms of run time, tangleGen and ADMIXTURE behave similarly on the Kidd’s AIMs panel, both needing about 20 seconds on a laptop with an AMD Ryzen 7 octa-core processor and 32 GB RAM. As the number of SNPs increases, both tangleGen and ADMIXTURE become significantly slower. However, tangleGen has the advantage that pre-computed costs can be saved and re-used to shorten runtime. A more detailed run time analysis is added in Supplemental Note 7.

### Data Simulation

We employ the software msprime (Baumdicker et al., 2022) to simulate the data with an underlying demographic structure for Fig. 4 A and Fig. 5 A. In the simulation for Fig. 4 A, migration has been added between populations A and E, population sizes are constant over time. We generated a total of 800 recombining diploid individuals characterized by 5041 biallelic SNPs. In the simulation for Fig. 5 A, the migration rate is quadrupled and added between all populations A to H. Again, we generated a total of 800 recombining diploid individuals and used the same mutation and recombination rate as in the simulation for Fig. 4 A, resulting in 5078 biallelic SNPs. The simulation scripts are available on Github at k-burger/tangles_in_pop_gen.

## Data Access

The data used in this study is available at Github (https://github.com/k-burger/tangles_in_pop_gen) for download or simulation. The code is also freely available on GitHub.

## Acknowledgements

This work has been supported by the Cluster of Excellence “Controlling Microbes to Fight Infections” (DFG, EXC 2124, Project number 390838134) and the Cluster of Excellence “Machine Lerning: New Perspectives for Science” (DFG, EXC 2064/1, Project number 390727645) and the International Max Planck Research School for Intelligent Systems (IMPRS-IS).

We acknowledge support by Open Access Publishing Fund of University of Tübingen.

