## Supplemental Material for "Inferring Ancestry with the Hierarchical Soft Clustering Approach tangleGen"

**1 ADMIXTURE Ancestry Proportions for the Minimal Example** **2**

**2 Results on Data from 1000 Genomes Project Including Ad-Mixed American Populations** **2**

**3 ADMIXTURE on 1000 Genomes Data Set up to $K = 7$** **3**

**4 Characteristic SNPs for Clustering of 1000 Genomes Data Set based on Kidd's AIMs Panel** **3**

**5 Cluster-Typical Set of SNPs for 1000 Genomes Data Set Based on Kidd's AIMs Panel** **3**

**6 tangleGen on Data with Many SNPs and Less Differentiated Populations** **4**

**7 Runtime Analysis** **7**

**8 FST Value Analysis for Kidd's AIMs Panel on 1000 Genomes Data** **8**

**9 fastStructure and SCOPE on Simulated and 1000 Genomes Data** **9**

**10 tangleGen on Simulated Data with More Migration** **10**

**11 Hyperparameters in tangleGen and their Effects** **11**

**12 Random Cost Function** **14**

**13 Effect of the Penalizing Factor $c_2$** **15**

#### Supplemental Note 1: ADMIXTURE Ancestry Proportions for the Minimal Example

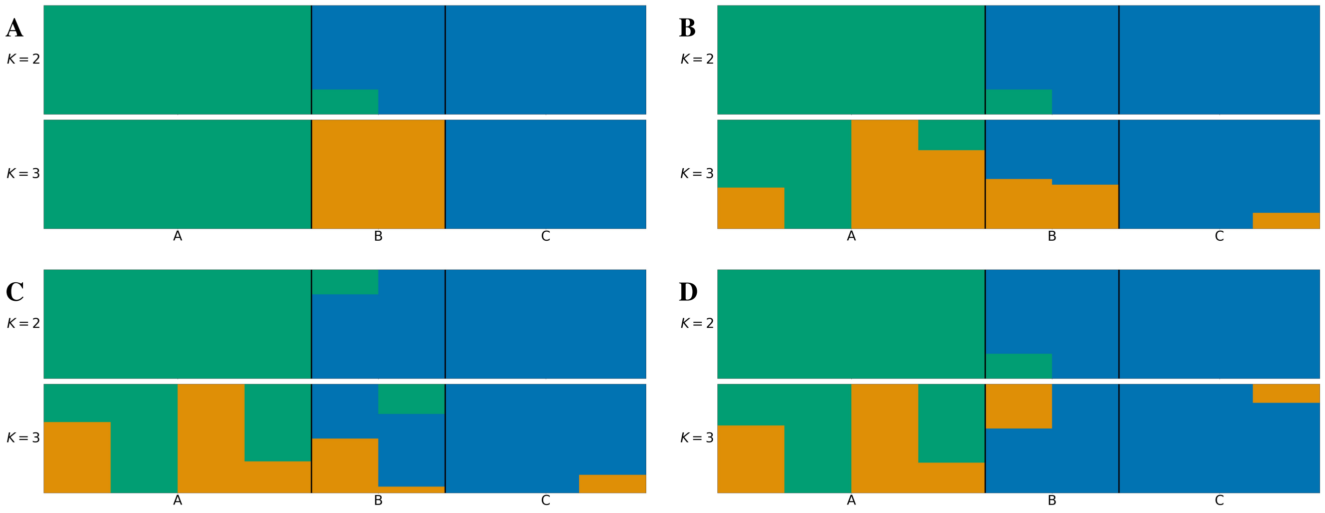


**Supplemental Fig. S1. A: ADMIXTURE ancestry proportions** on the minimal example from Results section. The best of ten runs is shown in terms of clarity of clusters and minimization of inconsistencies compared to the underlying population structure. The input parameter $K$, which denotes the number of populations to be identified, is specified for each subplot. ADMIXTURE identifies the three populations for $K = 3$. For this minimal exmaple, ADMIXTURE states three modes, the result in A is obtained in five out of ten runs. tangleGen's soft clustering on the same data set is shown in Fig. 3 B. **B, C & D: ADMIXTURE with different initial initializations.** For such a small data set, ADMIXTURE is particularly dependent on the initial initialization leading to significantly different estimates as shown in plots B, C and D. The ancestry proportions in B are achieved in three out of ten runs and in C and D in one out of ten runs each.

#### Supplemental Note 2: Results on Data from 1000 Genomes Project Including Ad-Mixed American Populations

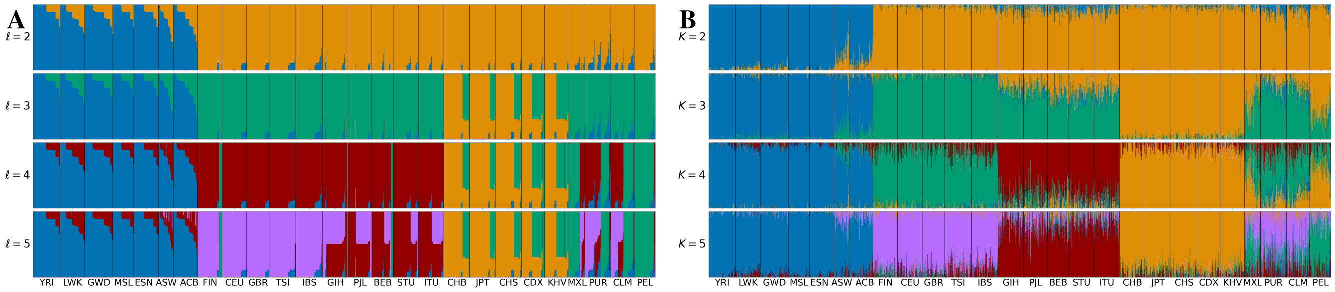


**Supplemental Fig. S2.** tangleGen and ADMIXTURE on the AIMs panel by Kidd et al. (2014) for individuals sampled in the data from 1000 Genomes Project Consortium et al. (2015) Phase 3, including Ad-Mixed American (AMR) populations. The individuals are sorted within the populations to achieve a block structure for tangleGen. The five superpopulations consist of the following populations: AFR: YRI, LWK, GWD, MSL, ESN, ASW, ACB; EUR: FIN, CEU, GBR, TSI, IBS; SAS: GIH, PJL, BEB, STU, ITU; EAS: CHB, JPT, CHS, CDX, KHV; AMR: MXL, PUR, CLM, PEL. **A: tangleGen soft clustering.** Each subplot corresponds to a level $\ell$ in the tangles tree, where the different levels result from splits in the tangles tree. tangleGen identifies the four superpopulations AFR, EUR, SAS and EAS, while AMR is composed of the other superpopulations. In Fig. 6, the EAS populations shared parts of their ancestry with the SAS populations and with the EUR populations. Here, no soft clustering between SAS and EAS is inferred, instead EAS now clusters partly with the Ad-mixed American populations. The first split in the tangles tree is supported by $9$ characteristic SNPs, the second by $2$, third by $1$ and forth by $2$. Parameters: Cost function as specified in Eq. 2, agreement parameter $a = 225$, $k = 40$. **B: ADMIXTURE ancestry proportions.** The best of ten ADMIXTURE runs is shown in terms of clarity of clusters and minimization of inconsistencies compared to the underlying population structure. The input parameter $K$, which denotes the number of populations to be identified, is specified for each subplot. ADMIXTURE identifies the four superpopulations AFR, EUR, SAS and EAS and even partially assigns AMR as a separate population. ADMIXTURE continues to show inconsistencies: for $K = 3$, for example, we observe that the South Asian populations share a considerable part of their ancestry with the East Asian populations (plotted in orange). However, for $K = 4$, this component disappears almost completely.

#### Supplemental Note 3: ADMIXTURE on 1000 Genomes Data Set up to $K = 7$

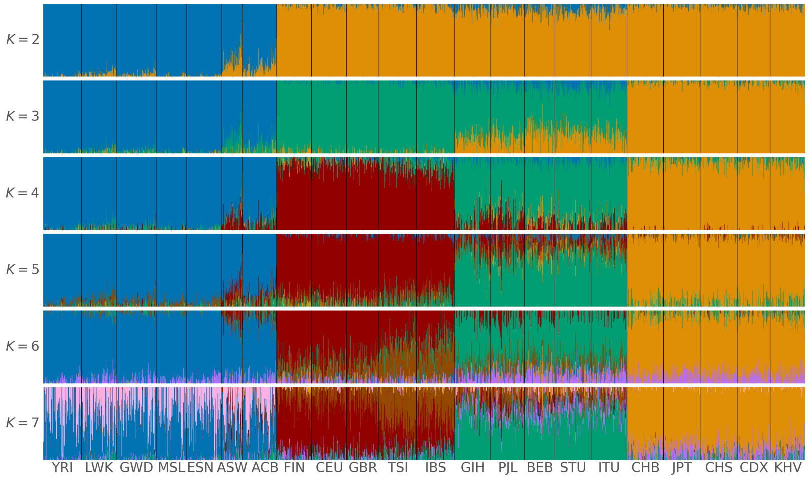


**Supplemental Fig. S3.** ADMIXTURE on Kidd's AIMs panel for individuals sampled in the 1000 Genomes Project Phase 3, excluding AMR populations. The four superpopulations are made up as follows: AFR: YRI, LWK, GWD, MSL, ESN, ASW, ACB; EUR: FIN, CEU, GBR, TSI, IBS; SAS: GIH, PJL, BEB, STU, ITU; EAS: CHB, JPT, CHS, CDX, KHV. The best of ten runs is shown in terms of clarity of clusters and minimization of inconsistencies compared to the underlying population structure. The input parameter $K$, which denotes the number of populations to be identified, is specified for each subplot. To compare ADMIXTURE with tangleGen, only $K$ values within the range considered by tangleGen are shown in Fig. 6 B in the main manuscript. This plot illustrates how ancestry proportions are estimated for larger $K$ values, that is $K > 4$. From $K = 5$, no further individual populations or superpopulations are identified; instead, superpopulations are shown as admixed.

#### Supplemental Note 4: Characteristic SNPs for Clustering of 1000 Genomes Data Set based on Kidd's AIMs Panel

A further advantage of the hierarchical clustering method tangleGen is that the SNPs responsible for the splits in the tangles tree and thus for the inferred ancestry can be identified. These SNPs are referred to as characteristic SNPs. Listed below are the characteristic SNPs for clustering individuals in the 1000 Genomes Project Phase 3 based on Kidd's AIMs panel as described in the Results section, Fig. 6, ordered from lowest to highest cost. We follow dbSNP ID notation by Sherry et al. (2001).

- *top split:* rs2814778, rs1871534, rs10497191, rs3916235, rs4891825, rs7326934, rs7554936, rs11652805, rs1462906.

- *second split:* rs3827760, rs1800414, rs3811801.

- *last split:* rs1426654, rs16891982, rs12913832.

#### Supplemental Note 5: Cluster-Typical Set of SNPs for 1000 Genomes Data Set Based on Kidd's AIMs Panel

tangleGen can extract a cluster-typical set of SNPs, comprising relevant cuts for each cluster found with the tangles tree alongside their orientations, enhancing the explainability of the method. A cluster-typical set of SNPs consists of all SNPs that were used for clustering, including those that were not characteristic of a particular split in the tangles tree. A cluster-typical set of SNPs currently indicates the presence of the derived allele without distinguishing between homozygous and heterozygous states as it is based on the underlying cuts, which cannot make a finer distinction due to the current design of cuts. Note that the cluster-typical set of SNPs does not represent the genome of any specific individual, as consistencies during the construction of the tangles tree are only ensured for triplets of cuts. In addition, the cluster-typical set of SNPs does not consist of all SNPs provided as input and usually varies in length for each cluster.

| SNP | AFR | EUR | SAS | EAS |
|---|---|---|---|---|
| rs2814778 | derived | ancestral | ancestral | ancestral |
| rs1871534 | derived | ancestral | ancestral | ancestral |
| rs3827760 | ancestral | ancestral | ancestral | derived |
| rs10497191 | ancestral | derived | derived | derived |
| rs3916235 | ancestral | derived | derived | derived |
| rs1800414 | ancestral | ancestral | ancestral | derived |
| rs4891825 | ancestral | derived | derived | derived |
| rs3811801 | ancestral | ancestral | ancestral | derived |
| rs1426654 | derived | ancestral | derived | derived |
| rs7326934 | derived | ancestral | ancestral | ancestral |
| rs7554936 | ancestral | derived | derived | derived |
| rs11652805 | ancestral | derived | derived | derived |
| rs16891982 | ancestral | derived | ancestral | ancestral |
| rs1462906 | ancestral | derived | derived | derived |
| rs1229984 | derived | derived | derived | |
| rs12913832 | ancestral | derived | ancestral | |
| rs1876482 | ancestral | ancestral | | |
| rs9522149 | ancestral | derived | | |
| rs310644 | derived | ancestral | | |
| rs3823159 | derived | ancestral | | |
| rs1834619 | ancestral | ancestral | | |
| rs798443 | ancestral | derived | | |
| rs7226659 | ancestral | ancestral | | |
| rs4918664 | ancestral | ancestral | | |
| rs2196051 | derived | | | |

**Supplemental Table S1.** Cluster-typical set of SNPs as a result of the clustering process by tangleGen for 1000 Genomes data set based on Kidd's AIMs panel. The resulting clusters can be associated with the superpopulations AFR, EUR, SAS, and EAS. In this context, "ancestral" denotes that the corresponding SNP is homozygous for the ancestral allele, while "derived" indicates that the corresponding SNP is homozygous or heterozygous for the derived allele. The characteristic SNPs within the tangles tree (Supplemental Note 4) are colored according to their corresponding split in the tangles tree, that is top split, second split and last split.

#### Supplemental Note 6: tangleGen on Data with Many SNPs and Less Differentiated Populations

This section discusses what happens when tangleGen is faced with two challenges simultaneously: first, that the underlying populations are not well differentiated and second, that the data contains many SNPs. It is clear that such a scenario poses a challenge for any hierarchical method, as considering too early a SNP positioned high in the ancestral tree may introduce population structures that are not representative of the actual data. By considering many SNPs, this risk increases considerably. Both these challenges are met, for example, when applying tangleGen to chromosome 22 in the 1000 Genomes Project data set. To provide a better understanding, we first examine simulated data in a similar setting. Moreover, for this scenario, a cost function based on the divergence of the Hardy-Weinberg equilibrium distinguishes the populations more clearly and is therefore introduced below.

**Cost Function Based on the Divergence from Hardy-Weinberg Equilibrium.** Let us use the same notation as in the Methods section and consider a cut $S_m$ that separates the individuals into subpopulations $A$ and $B = A^{\complement}$, $T = A \cup B$, with $N_A$, $N_B$ and $N$ the group sizes. In the following, we present a cost function that is also specific to population genetics, but targets different aspects compared to the FST-based cost function presented in the Methods section. While the FST-based cost function favors cuts that divide individuals into as different groups $A$ and $B$ as possible, we now want to use a cost function that favors cuts that lead to two well-mixed groups $A$ and $B$. To achieve this, we need to compare the expected heterozygosity with the observed heterozygosity in the groups. This evaluation metric is appropriate because these measures closely align in well-mixed subpopulations, that is a population in Hardy-Weinberg equilibrium. Depending on the application, a different cost function may be more suitable.

If the empirical frequency of the derived allele in group $A$ at SNP $S_m$ is given by

$$p_1^A = \frac{1}{2N_A} \sum_{n \in A} g_{nm}$$

then the expected heterozygosity in $A$ is

$$2p_1^A\left(1-p_1^A\right)$$

if $A$ is a well-mixed subpopulation. The number of observed heterozygotes in group $A$ at SNP $S_m$ is

$$x_{01}^A = \frac{1}{N_A}\sum_{n\in A}\mathbb{1}_{g_{nm}=1}$$

and therefore, the mean divergence from the Hardy-Weinberg equilibrium over all SNPs is given by

$$\bar{D}^A = \frac{1}{M}\sum_{m=1}^{M}|x_{01}^A - 2p_1^A(1-p_1^A)|.$$

Finally, the cost function is given by

$$c_{\text{HWE}}(S_m) := \frac{c_1^2\cdot c_2}{c_3}\cdot\left(1+\min\left(\bar{D}^A,\bar{D}^B\right)\right) \tag{1}$$

with

$$c_1 = 1+\left(\frac{\text{kNN\_A\_in\_B}}{N_A\cdot 40}+\frac{\text{kNN\_B\_in\_A}}{N_B\cdot 40}\right)$$

and

$$c_2 = \left(\frac{N}{N_A}+\frac{N}{N_B}\right)^{0.1}$$

as introduced in Eq. 3 and Eq. 4 in the main manuscript and

$$c_3 = \frac{1}{M}\sum_{m=1}^{M}|p_1^A - p_1^B|^{\frac{1}{2}}$$

a penalization factor that accounts for the differences between $A$ and $B$, not just the differences within the groups. This also ensures that random cuts through an overall population that is in Hardy-Weinberg equilibrium does not receive a very low cost leading to a prioritization by tangleGen.

***Comparison with the Hardy-Weinberg-Equilibrium based reliability factor in the soft clustering step.*** When using this cost function, both the cost function and soft clustering are based on the Hardy-Weinberg equilibrium, which might suggest that they have the same effect. However, this is not the case. The soft clustering works with individual SNPs and indicates their reliability in population differentiation with respect to Hardy-Weinberg equilibrium. Conversely, the cost function evaluates each SNP/cut for its division into two groups based on all other SNPs and assigns a low cost if the two groups are significantly different and well-mixed. This is also evident in the minimal example, where the Hardy-Weinberg-based cost function assigns the same and lowest cost to both SNPs $s_3$ and $s_6$, while SNP $s_5$ has a higher cost. In contrast, only $s_5$ and $s_6$ are given a reliability factor of 1 by the soft clustering, as they are both homozygous for every individual.

**Simulation.** In the following simulation, we used the same simulation script with the same underlying demographic structure as previously described in Fig. 4 A. However, we reduced the time between population splits by a factor of 10 to obtain less differentiated populations. In addition, we increased the mutation rate by a factor of 20 to generate significantly more SNPs. After deleting SNPs with an allele frequency of less than 5% or higher than 95%, a total of 9420 SNPs are considered for the clustering of 800 individuals. The simulation script is available on Github at k-burger/tangles_in_pop_gen.

**Comparison of two Cost Functions on Simulated Data with Many SNPs and Less Differentiated Populations.** To compare the FST-based cost function introduced in the main manuscript with the cost function stated in in this Supplemental Note, we apply tangleGen twice to the simulated data with many SNPs and less differentiated populations from supplemental simulation section. Fig. S4 A shows that the FST-based cost function has difficulties in this scenario compared to Fig. 4 D, especially since it is not able to distinguish between the populations A and B as well as E and F, that is the populations that have split last in the underlying population structure (Fig. 4 A) and are therefore not well-differentiated. In contrast, tangleGen with the Hardy-Weinberg-based cost function, Fig. S4 B, shows a clearer clustering compared to Fig. S4 A. However, this cost function does not reveal the underlying population structure when applied to the simulated data described in Fig. 4 A, as it tends to split the most differentiated individual populations first (like population $H$).

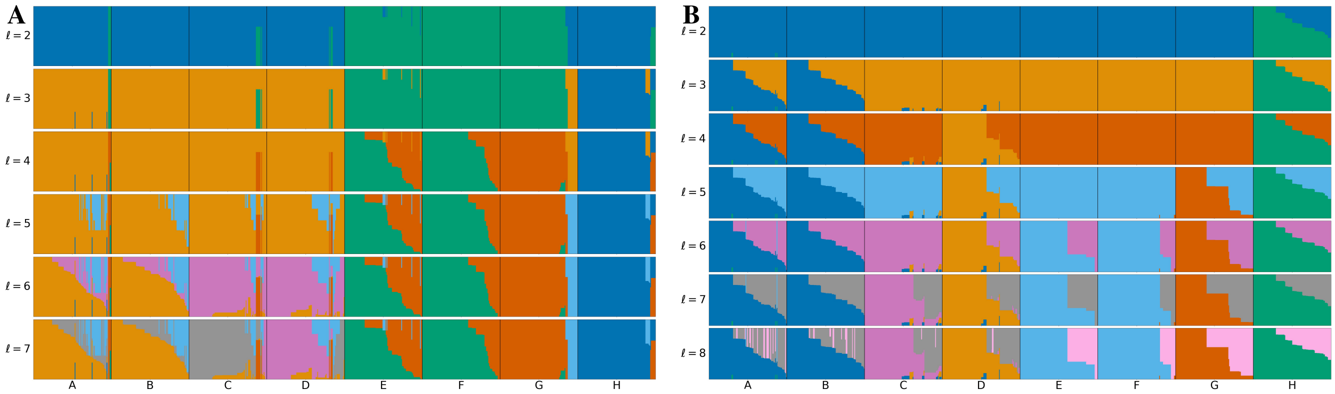


**Supplemental Fig. S4.** tangleGen on simulated data with many SNPs and less differentiated populations as described in the supplemental simulation section and two different cost functions. The individuals are sorted within the populations to achieve a block structure. Each subplot corresponds to a level $\ell$ in the tangles tree, where the different levels result from splits in the tangles tree. **A: tangleGen with FST-based cost function.** Used cost function as in Eq. 2, agreement parameter $a = 40$, pruning is set to $2$ and $k = 40$. tangleGen clusters the individuals hierarchically into seven populations. However, tangleGen does not distinguish between the most closely related populations A and B and E and F, the populations that split last in the underlying population structure (Fig. 4 A) and are therefore not well differentiated. Instead tangleGen identifies clusters that correspond rather to ancestral populations instead of extant populations. For example parts of population E and F cluster together with population G. Starting with the top split between the ancestral populations ABCDH and EFG, each population split is based on $15$, $12$, $26$, $16$, $25$ and $3$ characteristic SNPs. **B: tangleGen with Hardy-Weinberg-based cost function.** Used cost function as in Eq. 1, agreement parameter $a = 40$, pruning is set to $2$ and $k = 40$. tangleGen with this cost function is similar to A but results at level $\ell = 8$ in a clearer distinction and explicit separation of left and right subtree within the underlying population structure from Fig. 4, i.e. between the ancestral populations ABCD (pink cluster) and EFGH (gray cluster). Furthermore, the order of splits varied due to the different cost function. Starting with the top split between the ancestral populations ABCDEFG and H, each population split is based on $47$, $11$, $8$, $7$, $3$, $5$ and $4$ characteristic SNPs.

In contrast to tangleGen, the scenario with less differentiated populations and more SNPs is much less of a challenge for ADMIXTURE, as Fig. S5 shows for $K = 8$. This outcome is not surprising as ADMIXTURE benefits from having many SNPs available and does not face the challenges of an hierarchical method. However, ADMIXTURE still performs significantly worse than in Fig. 4 and also displays inconsistencies with the underlying population structure.

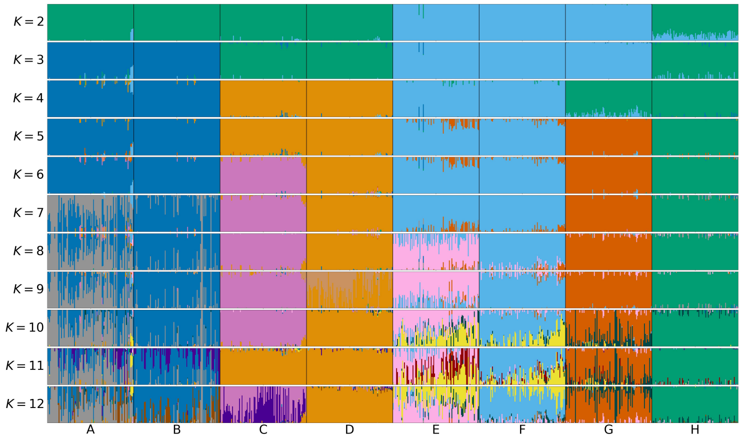


**Supplemental Fig. S5.** ADMIXTURE on simulated data with many SNPs and less differentiated populations as described in the supplemental simulation section. The best of ten runs is shown in terms of clarity of clusters and minimization of inconsistencies compared to the underlying population structure. The individuals are sorted within the populations in the same way such that a block structure for the tangleGen in Fig. S4 is achieved. The input parameter $K$, which denotes the number of populations to be identified, is specified for each subplot. For $K = 8$ ADMIXTURE separates the eight populations comparably well to Fig. S4. But in this scenario the separation is less clear compared to the scenario in Fig. 4 and again shows inconsistencies for example between $K = 3$ and $K = 4$. For $K \geq 9$, the further inferred admixtures do and cannot represent meaningful parts of the underlying demography with eight populations.

**Results on full 1000 Genomes Data Set.** The application of tangleGen with the Hardy-Weinberg-based cost function on chromosome 22 (from the 1000 Genomes Project, Phase 3 data set) is shown in Fig. S6. At first glance, the clustering appears similar to Fig. 6 B, which is based on Kidd's AIMs panel. However, the differentiation of superpopulations is less pronounced, especially at the lowest level, when European and South Asian populations are distinguished. This observation is further reinforced when considering the number of SNPs responsible for each split in the tangles tree. While the Hardy-Weinberg-based cost function performs better than the FST-based one in this scenario, further fine-tuning of the cuts and cost functions might be able to address the inherent challenges of hierarchical methods in such scenarios.

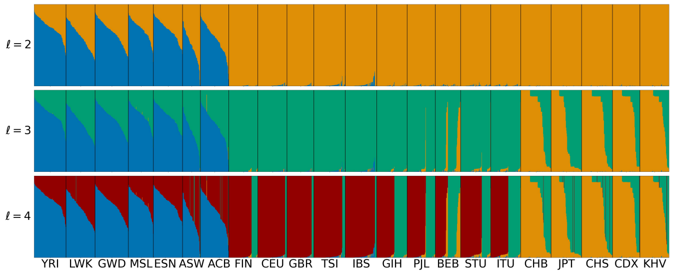


**Supplemental Fig. S6.** tangleGen on chromosome $22$ (1000 Genomes Project, Phase $3$), AMR populations are excluded and SNPs in frequncy below 5% are ignored. The individuals are sorted within the populations to achieve a block structure for tangleGen. Each subplot corresponds to a level $\ell$ in the tangles tree, where the different levels result from splits in the tangles tree. Hardy-Weinberg-based cost function is used (as stated in Eq. 1), agreement parameter $a = 200$, no pruning added and $k = 40$. tangleGen identifies the African and East Asian populations but has difficulties in differentiating European and South Asian populations. The first split is supported by $181$ characteristic SNPs, the second by $37$ and the third by only a single cut. If the pruning option in tangleGen is used, the last split would thus be omitted.

### Supplemental Note 7: Runtime Analysis

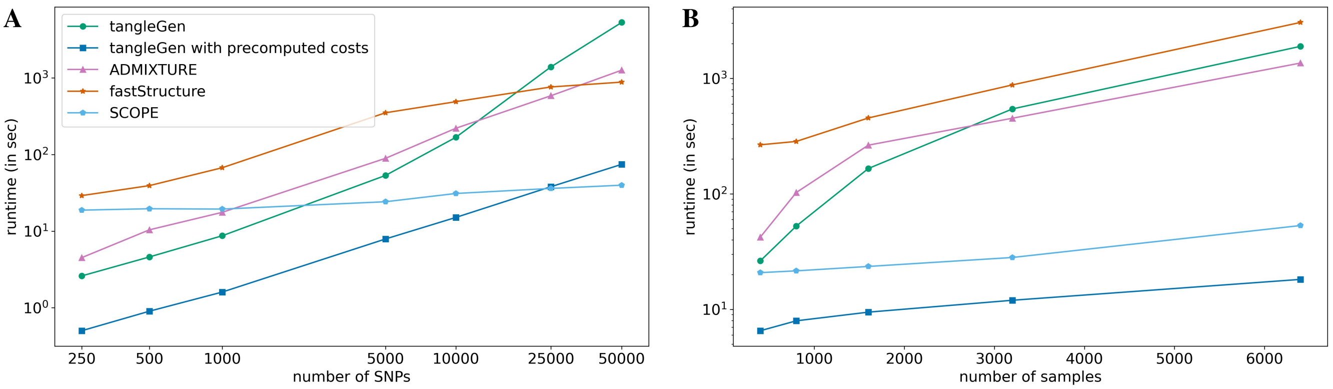


**Supplemental Fig. S7.** The runtime analysis compares tangleGen with ADMIXTURE, fastStructure (Raj et al., 2014) and SCOPE (Chiu et al., 2022) using simulated data as described in Fig. 4 A and the Data Simulation section in Methods. tangleGen uses the FST-based cost function with hyperparameters $k = 40$ and $b = 0.05$. The maximum input parameter $K$ (eight in this analysis) for ADMIXTURE, fastStructure and SCOPE is determined by the maximum number of populations identified by tangleGen. It is important to note that determining this $K$ value in practice requires time and careful consideration, which has not been factored in here. The calculations were performed on a laptop with an AMD Ryzen 7 Octa-Core processor and 32 GB RAM. **A: Number of SNPs.** We observe that tangleGen performs faster than ADMIXTURE and fastStructure on smaller data sets. However, as the size of the data set increases, tangleGen becomes significantly slower, which is due to the cost and tangles tree computation. As the cost calculation precedes the actual Tangles algorithm, it is a process that can be saved and reloaded for subsequent runs, for example when adjusting the agreement parameter. The calculation of the tangles tree is computationally challenging on large datasets, as for each SNP to be added, the consistency with all triplets of SNPs already added is checked for each branch. To circumvent this and improve scalability, one could subsample the triplets above a certain SNP count instead of considering all of them. This will improve the runtime without noticeably affecting performance and will be part of future research. SCOPE scales very well to larger data. Number of samples in this analysis: $800$. **B: Number of samples.** Increasing the number of samples increases the runtime of all methods considered. As sample sizes grow, the cost calculation with tangleGen becomes increasingly time-consuming. However, with precalculated costs, tangleGen has the lowest runtime for all sample sizes considered. Number of SNPs in this analysis: $5041$.

#### Supplemental Note 8: FST Value Analysis for Kidd's AIMs Panel on 1000 Genomes Data

| split-off superpopulation | SNP | FST value | characteristic SNP usage rank | cost |
|---|---|---|---|---|
| AFR | rs2814778 | 0.94 | 1 | 2.35 |
| | rs1871534 | 0.87 | 2 | 2.38 |
| | rs3916235 | 0.69 | 4 | 2.84 |
| | rs7326934 | 0.65 | 6 | 3.17 |
| | rs4891825 | 0.64 | 5 | 2.87 |
| | rs3823159 | 0.62 | - | 4.14 |
| | rs10497191 | 0.62 | 3 | 2.78 |
| | rs1462906 | 0.58 | 9 | 3.53 |
| | rs7657799 | 0.51 | - | 4.16 |
| | rs7251928 | 0.50 | - | 4.64 |
| | rs310644 | 0.48 | - | 4.13 |
| | rs7554936 | 0.47 | 7 | 3.25 |
| | rs2593595 | 0.45 | - | 5.69 |
| | rs11652805 | 0.45 | 8 | 3.29 |
| EAS | rs3827760 | 0.81 | 1 | 2.63 |
| | rs1229984 | 0.58 | - | 3.85 |
| | rs1800414 | 0.52 | 2 | 2.85 |
| | rs1876482 | 0.45 | - | 4.05 |
| | rs3811801 | 0.44 | 3 | 3.00 |
| EUR | rs16891982 | 0.79 | 2 | 3.38 |
| | rs12913832 | 0.45 | 3 | 3.92 |
| | rs9522149 | 0.44 | - | 4.12 |
| | rs1426654 | 0.43 | 1 | 3.07 |

**Supplemental Table S2.** FST value analysis for Kidd's AIMs panel on 1000 Genomes Data. This analysis considers three splits: AFR vs. EUR/SAS/EAS, EAS vs. AFR/EUR/SAS and EUR vs. AFR/SAS/EAS. For each split, only the SNPs with the highest FST values are listed, sorted by FST value. tangleGen identifies 9 characteristic SNPs for the first split and three each for the second and third split. For each SNP, it is indicated whether it was identified by tangleGen as a characteristic SNP and, if so, the usage rank of this SNP (SNP with rank 1 has the greatest impact, as it is considered first by the hierarchical method). The cost assigned by tangleGen is also given. Since tangleGen assigns each SNP a cost based on the mean FST value over all SNPs with respect to the division of individuals induced by the corresponding cut, there is a correlation between the characteristic SNPs and the SNPs with the highest FST values. In addition, the cost function takes k-nearest neighbors and the balance of bipartition sizes into account. Furthermore, the calculation of the true FST values is based on the classification into superpopulations, information that is not available for tangleGen. Some SNPs have a high FST value but are not characteristic SNPs. This is because they have a significantly higher cost and tangleGen composes the tangles tree by orienting cuts/SNPs iteratively beginning with the lowest-cost cut. The tangles tree is complete when no further cuts can be consistently added. From this moment any SNP with a higher cost is no longer considered. In a deeper tangles tree (for example with a smaller agreement parameter), some high FST SNPs might be considered characteristic (for example rs1229984 for the EAS split), but they would have a lower rank than the characteristic SNPs in this analysis.

#### Supplemental Note 9: fastStructure and SCOPE on Simulated and 1000 Genomes Data

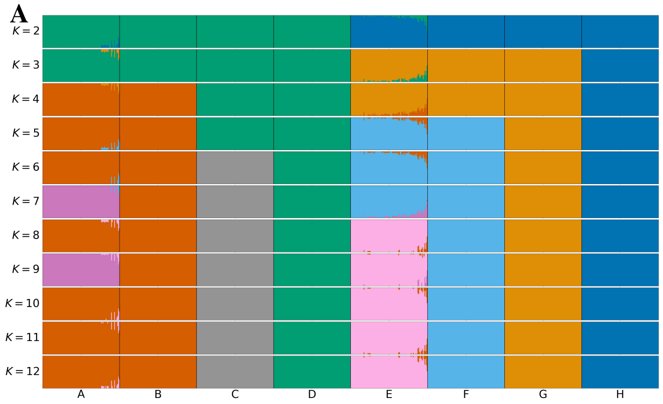


**Supplemental Fig. S8.** Inferred ancestries by fastStructure and SCOPE from simulated genetic data as in Fig. 4 A for different pre-specified numbers of populations, denoted by $K$. Individuals are sorted within the populations in the same order as in the tangleGen soft custer plot Fig. 4 D. The best of ten runs is shown in terms of clarity of clusters and minimization of inconsistencies compared to the underlying population structure. **A: fastStructure.** fastStructure estimates the ancestry proportions well for $K \leq 8$ but has difficulties to seperate populations A and B. The number of identified populations does not match $K$ for $K > 7$ since fastStructure can generate empty populations and the simulated data does not contain further population structure for $K > 8$. Furthermore, the results for different $K$ show hierarchical inconsistencies, particularly for $K \geq 8$ in populations A and B. **B: SCOPE.** SCOPE estimates ancestry proportions well for the true number of populations ($K = 8$) and provides plausible results for other values of $K$. For $K > 8$, the simulated data does not contain additional population structure. The non-hierarchical nature of SCOPE is evident, for example, in population H between $K = 3$ and $K = 4$.

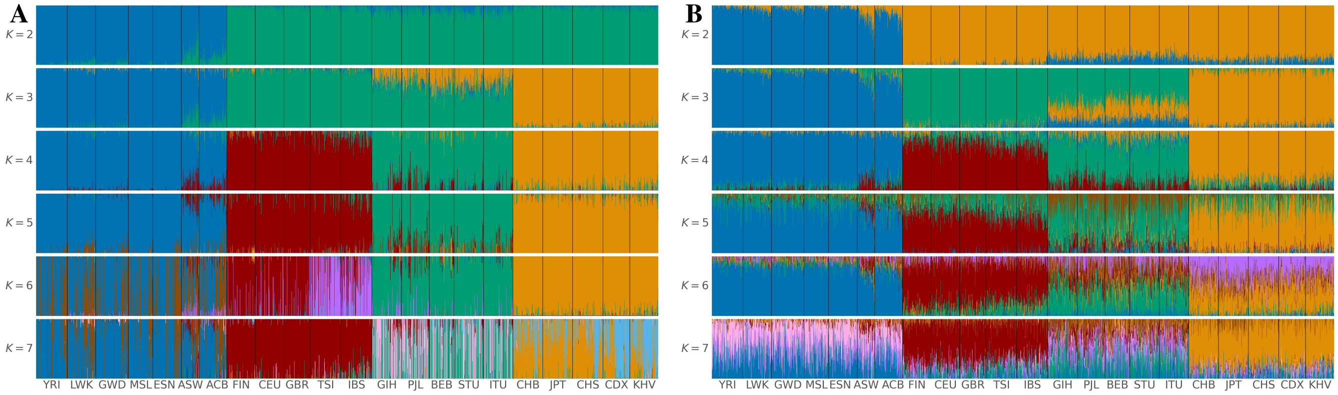


**Supplemental Fig. S9.** fastStructure and SCOPE on Kidd's AIMs panel for individuals sampled in the 1000 Genomes Project Phase 3, excluding AMR populations. The four superpopulations are made up as follows: AFR: YRI, LWK, GWD, MSL, ESN, ASW, ACB; EUR: FIN, CEU, GBR, TSI, IBS; SAS: GIH, PJL, BEB, STU, ITU; EAS: CHB, JPT, CHS, CDX, KHV. The best of ten runs is shown in terms of clarity of clusters and minimization of inconsistencies compared to the underlying population structure. For each subplot the input parameter $K$, which denotes the pre-specified number of independent populations to be identified, is shown on the left side. fastStructure **(A)** and SCOPE **(B)** identify the four superpopulations AFR, EUR, SAS and EAS, when the number of populations is set to $K = 4$, but show inconsistencies, compared to smaller values of $K$ in the differentiation between the four populations (for example between $K = 3$ and $K = 4$). Interestingly, fastStructure recognizes TSI and IBS as a separate population for $K = 6$ although Kidd's AIMs panel was not optimized for this. Beyond this, superpopulations are presented as admixed for $K > 4$.

#### Supplemental Note 10: tangleGen on Simulated Data with More Migration

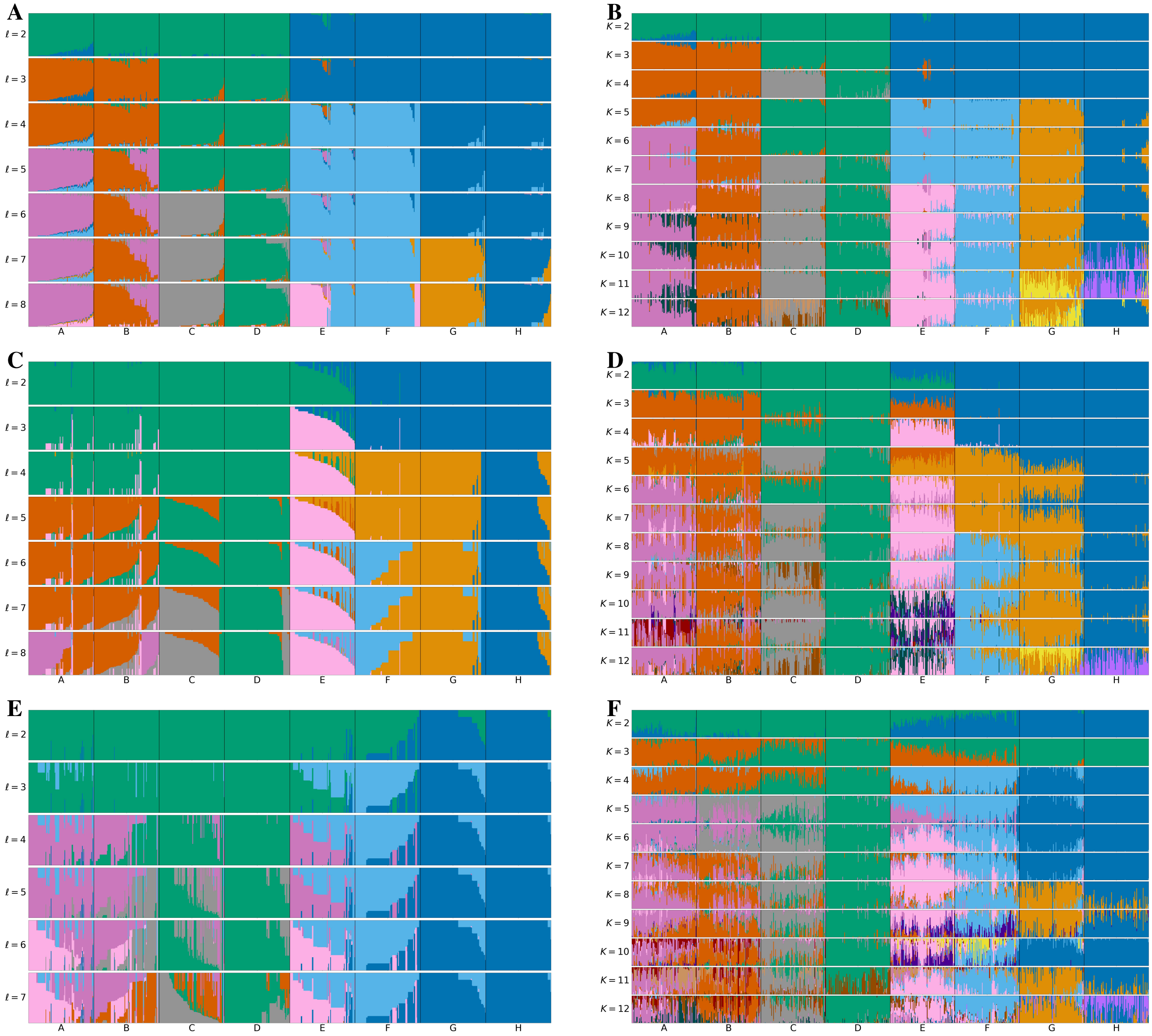

**Supplemental Fig. S10. Comparison of tangleGen and ADMIXTURE on simulated data, as described in Fig. 5 A, but with varying migration rates between populations A to H**. The first column (A, C, and E) shows tangleGen's soft clustering visualized in bar plots for different level $\ell$ in the tangles tree. The individuals are sorted within the populations based on the soft cluster proportions. For this analysis, the agreement parameter is set to $a = 30$ and all external branches supported by only one SNP are pruned. The second column (B, D, and F) shows ADMIXTURE's ancestry proportions for different pre-specified numbers of populations ($K$). Individuals are sorted within populations in the same order as in the corresponding tangleGen plot. In contrast to tangleGen, ADMIXTURE is not deterministic and the best of ten runs is shown in terms of clarity of clusters and minimization of inconsistencies compared to the underlying population structure. Migration rates increase from top to bottom: In **A** and **B**, the migration rate is as in Fig. 4 between populations A and E. In subplots **C** and **D**, the rate is quadrupled (as in Fig. 5); in **E** and **F** doubled again. **A, C** and **E** show that tangleGen infers meaningful population structures even when increased migration leads to more admixed populations. The hierarchical nature of tangleGen helps to clarify the migration pattern. For the highest migration rate (**E**) tangleGen cannot differentiate G and H, which is not surprising as these populations split early in the underlying population structure and thus experience the most migration. Furthermore, increased migration reduces the number of SNPs that separate populations well. Consequently, tangleGen's clustering is based on fewer SNPs. Starting with the top split between the ancestral populations each split in A is based on 94 (C:29, E:8), 100 (C:11, E:6), 14 (C:10, E:9), 14 (C:36, E:6), 16 (C:10, E:8), 17 (C:4, E:8) and 2 (C:3) characteristic SNPs. Subplots **B, D**, and **F** show ADMIXTURE's performance, estimating populations well for the true number of populations ($K = 8$). While the migration patterns are also visible with ADMIXTURE, they are not as clear as with tangleGen due to ADMIXTURE's non-hierarchical approach (A and C). ADMIXTURE results for different $K$ show inconsistencies in each plot (**B, D**, and **F**). For $K > 8$, the simulated data does not contain any further population structure. Unlike ADMIXTURE, which requires choosing $K$, tangleGen determines the number of populations as a result of its hierarchical clustering in combination with the agreement parameter at $\ell = 8$ or $\ell = 7$.

#### Supplemental Note 11: Hyperparameters in tangleGen and their Effects

tangleGen uses several hyperparameters: the agreement parameter $a$, the pruning parameter and two additional parameters $k$ and $b$ in the penalization factors of the FST-based cost function. Here, $k$ is related to the $k$-nearest neighbors (kNN) factor $c_1$, and $b$ controls the preference for balanced cuts in $c_2$. To assess the impact of these hyperparameters on tangleGen's clustering results, we use a modified version of Dasgupta's hierarchical quality measure (Dasgupta, 2016)

$$D = \sum_{i,j} w(i,j) \cdot |C(i,j)|$$

where $w(i,j)$ denotes the pairwise difference between individuals $i$ and $j$ and $|C(i,j)|$ represents the size of the smallest cluster containing both individuals. It is important to note that Dasgupta's measure evaluates hard clustering, where each individual is assigned exactly one cluster, unlike the soft clustering shown in the previous plots. However, the hard clustering can be computed from the soft clustering as discussed in the Methods section. Furthermore, Dasgupta's measure assumes clustering down to the individual level, which differs from tangleGen approach of clustering individuals into populations. Consequently, the measure does not detect overclustering, as smaller cluster sizes decrease $D$. This limitation is mitigated to some extent by setting $w(i,j) = 0$ for individuals sampled from the same population. To detect overclustering, we consider the silhouette score (Rousseeuw, 1987), which effectively identifies small, random clusters but tends to overlook underclustering, a strength of Dasgupta's measure.

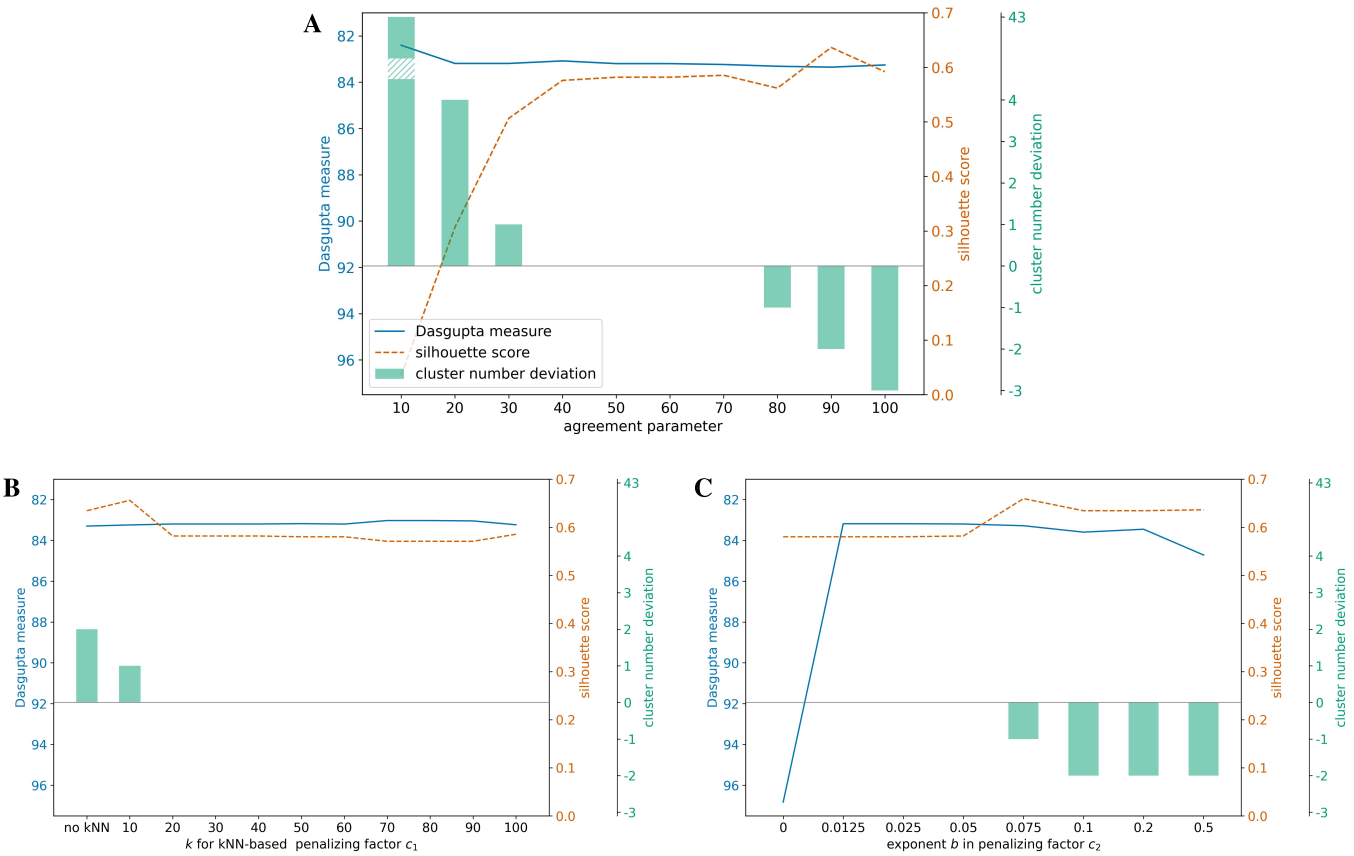


**Supplemental Fig. S11. Hyperparameter effects on simulation**, as in Fig. 4 A, with constant population sizes of $100$. The following analysis is based on hard clustering, with pruning omitted since it only affects boundary values. **A: Agreement parameter.** tangleGen performs best with intermediate agreement parameters, yielding good values in both Dasgupta's measure and silhouette score. Low agreement parameters lead to significant overclustering, shown by cluster number deviation, and low silhouette scores. Since Dasgupta's measure is not sensitive to overclustering, it assigns a small error in such cases. Conversely, high agreement parameters cause underclustering, resulting in higher silhouette scores, as this measure tends to overlook underclustering. Dasgupta's measure shows slightly higher errors than for intermediate agreement parameters. These observations align with tangles' recommendations (Klepper et al., 2023). Other hyperparameters: $k = 40,\ b = 0.05$. **B: $k$ for kNN-based penalizing factor $c_1$.** Adding a kNN-based penalizing factor to the cost function is beneficial, with the choice of $k$ being relatively insensitive as long as it is not too small. Other hyperparameters: $a = 50,\ b = 0.05$. **C: Exponent $b$ in penalizing factor $c_2$.** Adding a penalizing factor that prefers balanced cuts benefits the hierarchical method tangleGen. However, this parameter is sensitive; a too large $b$ reduces cluster quality by assigning too low a cost to unfavorable clusters, as shown in Fig. S16. Other hyperparameters: $a = 50,\ k = 40$.

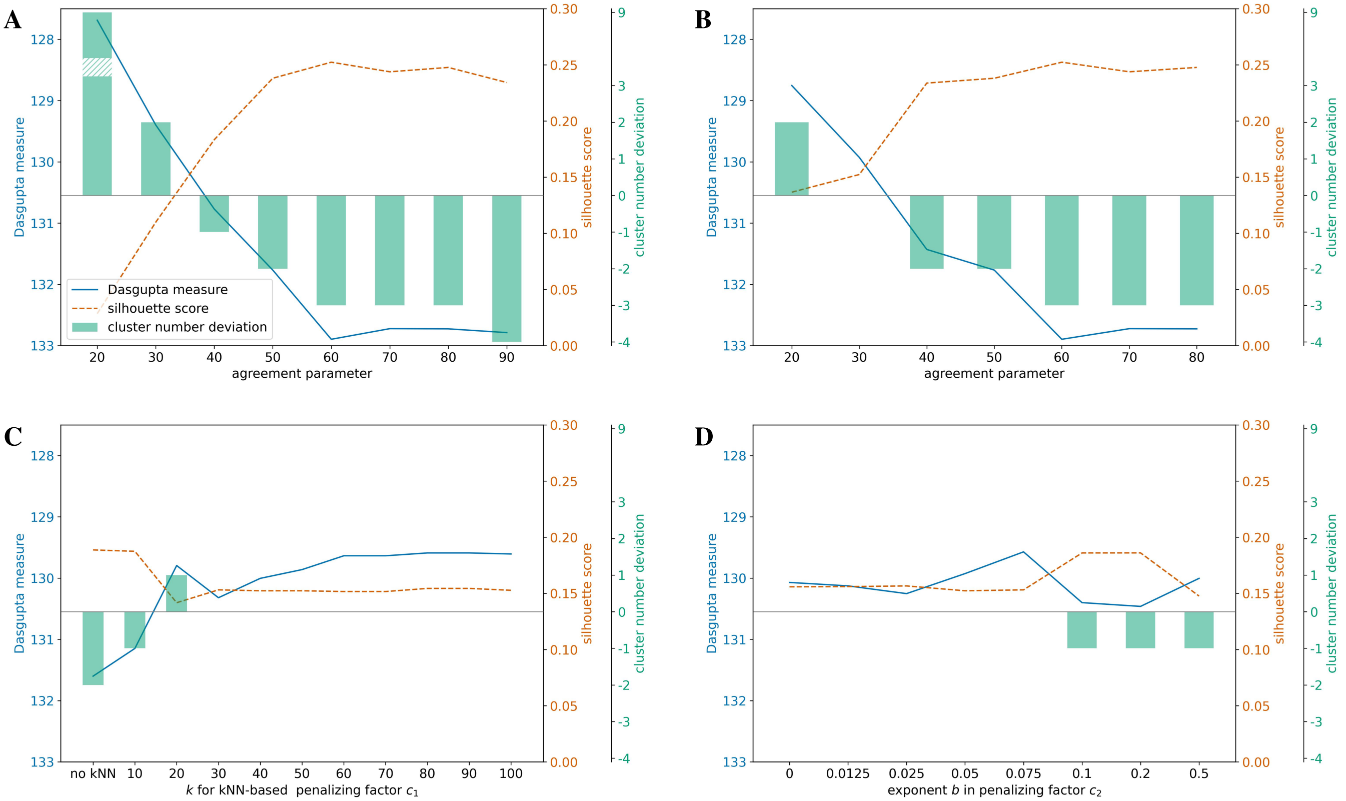


**Supplemental Fig. S12. Hyperparameter effects on simulation with migration between populations A to H**, as in Fig. 5 A, with constant population sizes of $100$. The following analysis is based on hard clustering. **A: Agreement parameter (no pruning).** This complex simulation is more challenging for tangleGen than the scenario in Fig. S11 A. This difficulty is reflected in the generally higher values for the Dasgupta measure and the lower silhouette scores. The agreement parameter is particularly sensitive in this context. Unlike in Fig. S11 A, there is no section where both measures indicate a high quality. Other hyperparameters: $k = 40,\ b = 0.05$. **B: Agreement parameter (external branches of length two pruned).** With pruning clustering becomes more stable in this scenario. Other hyperparameters: $k = 40,\ b = 0.05$. **C: $k$ for kNN-based penalizing factor $c_1$.** As observed in Fig. S11 B, adding a kNN-based penalty factor to the cost function is beneficial, and the choice of $k$ is relatively insensitive as long as it is not too small. Other hyperparameters: $a = 30,\ b = 0.05$, external branches of length two pruned. **D: Exponent $b$ in penalizing factor $c_2$.** Using a penalization factor that favors balanced cuts does not have as great an effect in this scenario as in Fig. S11 C. As before, if $b$ is chosen too large, tangleGen's clustering is negatively affected. Other hyperparameters: $a = 30,\ k = 40$, external branches of length two pruned.

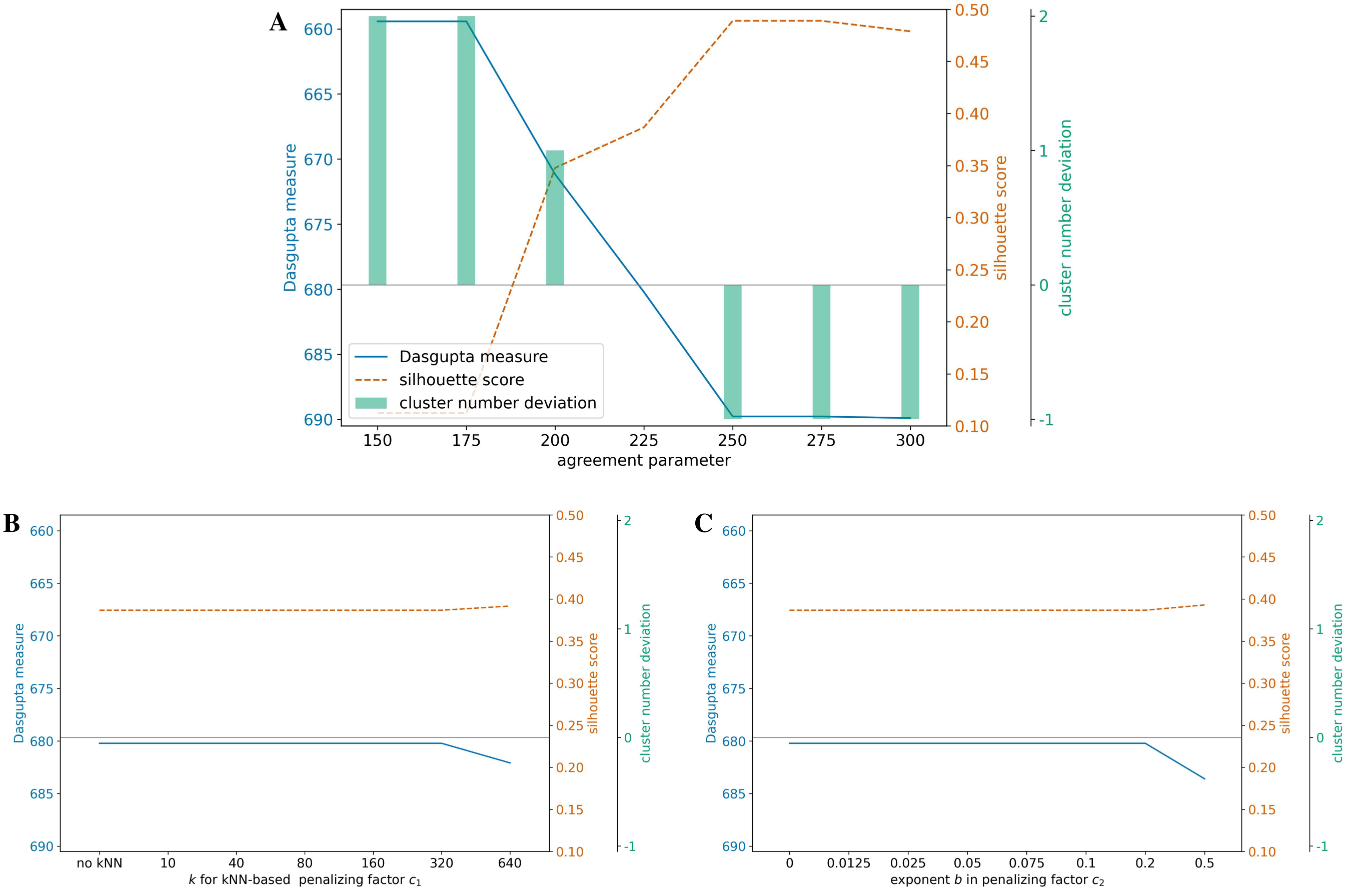


**Supplemental Fig. S13. Hyperparameter effects on 1000 Genomes data** (Fig. 6). The following analysis is based on hard clustering, with pruning disabled. **A: Agreement parameter.** The agreement parameter is a sensitive hyperparameter in this context. tangleGen's clustering shows overclustering when the parameter is set too low and underclustering when it is set too high, as indicated by the two quality measures and the cluster number deviation. tangleGen performs best with intermediate agreement parameters ($225$ corresponds to about 34% of the largest superpopulation (AFR) and 46% of the smallest superpopulation (SAS)), aligning with tangles' recommendations (Klepper et al., 2023). Other hyperparameters: $k = 40,\ b = 0.05$. **B & C: $k$ for kNN-based penalizing factor $c_1$ and exponent $b$ in penalizing factor $c_2$.** The penalization factors $c_1$ and $c_2$ are added to the cost function to enhance stability against cuts separating closely related individuals and to prevent the early onset of inconsistencies caused by unbalanced cuts. In this context, however, we consider Kidd's AIMs panel providing a stable data basis. As a result, $c_1$ and $c_2$ do not have a significant effect on this dataset as long as $k$ and $b$ are not chosen too large. Other hyperparameters: $a = 225,\ b = 0.05$ in A, $a = 225,\ k = 40$ in B.

#### Supplemental Note 12: Random Cost Function

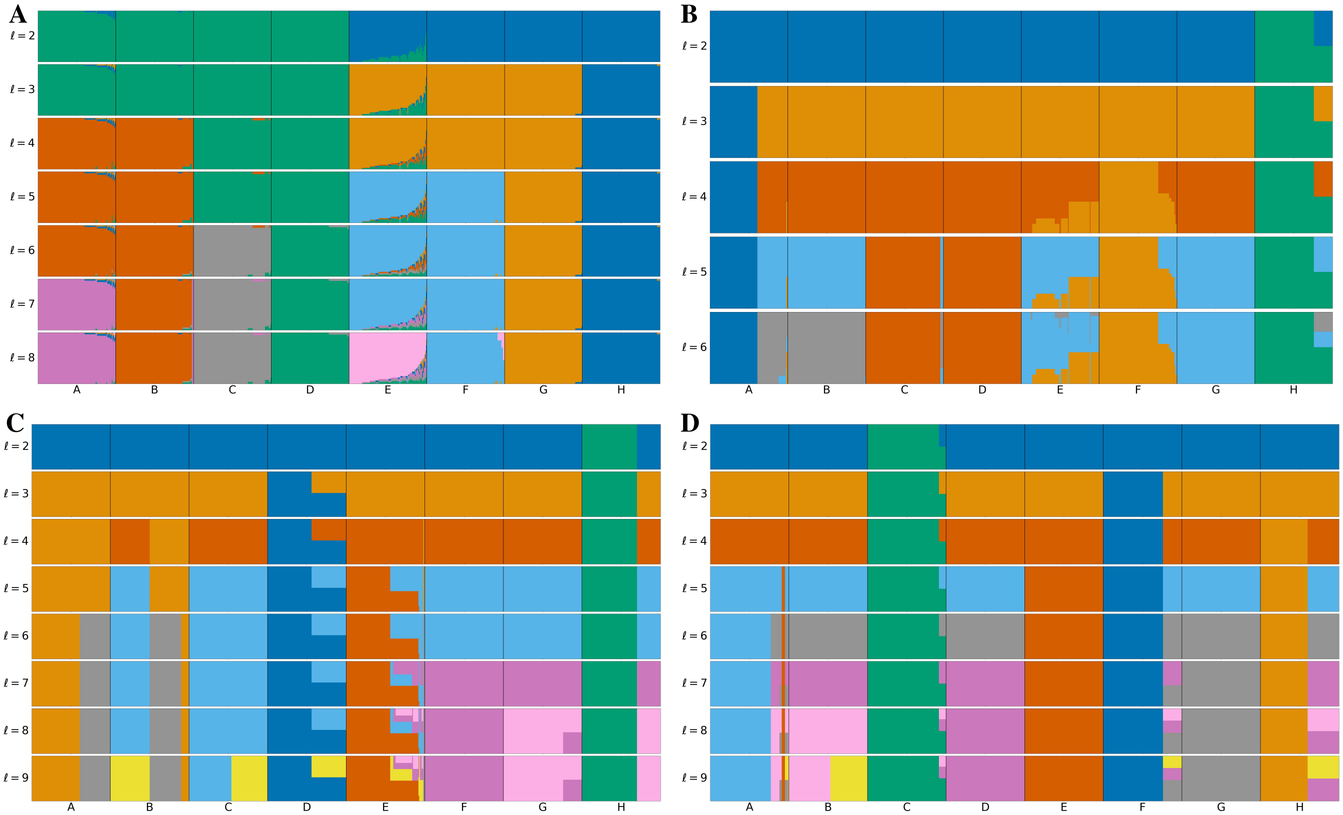


**Supplemental Fig. S14. Random cost function on simulated data.** The cost function has a significant impact on the performance of tangleGen. Based on the cost function, the cuts are sorted and iteratively oriented to compose the tangles tree. This analysis shows the effects of assigning random costs to each SNP/cut using the simulated data from Fig. 4 A. Panel **A** shows tangleGen with the FST-based cost function for ease of comparison. Panels **B**, **C**, and **D** show tangleGen with random cost assignments using different seeds. The individuals are sorted within the populations based on the soft cluster proportions to achieve a block structure in the plots. Each subplot corresponds to a level $\ell$ in the tangles tree, where the different levels result from splits in the tangles tree. The agreement parameter is set to $a = 50$ in all panels. Without a meaningful cost function, both population splits and the order of splits are no longer consistent with the underlying population structure. In particular, tangleGen almost always splits off individual populations instead of recognizing larger clusters, such as ABCD vs. EFGH for $\ell = 2$. Moreover, the depth of the inferred hierarchy varies depending on the SNPs used first, and the inference is based on significantly fewer SNPs per level than before. Nevertheless, tangleGen succeeds in identifying some individual populations, for example population F in panel C and populations D, E, and G in panel D for $\ell = 9$. This is due to the clear differentiation between the populations in this simulation, leading to a significant amount of SNPs with high discriminatory power to which low costs can be randomly assigned. This is no longer the case for data with less differentiation, as shown in Fig. S15.

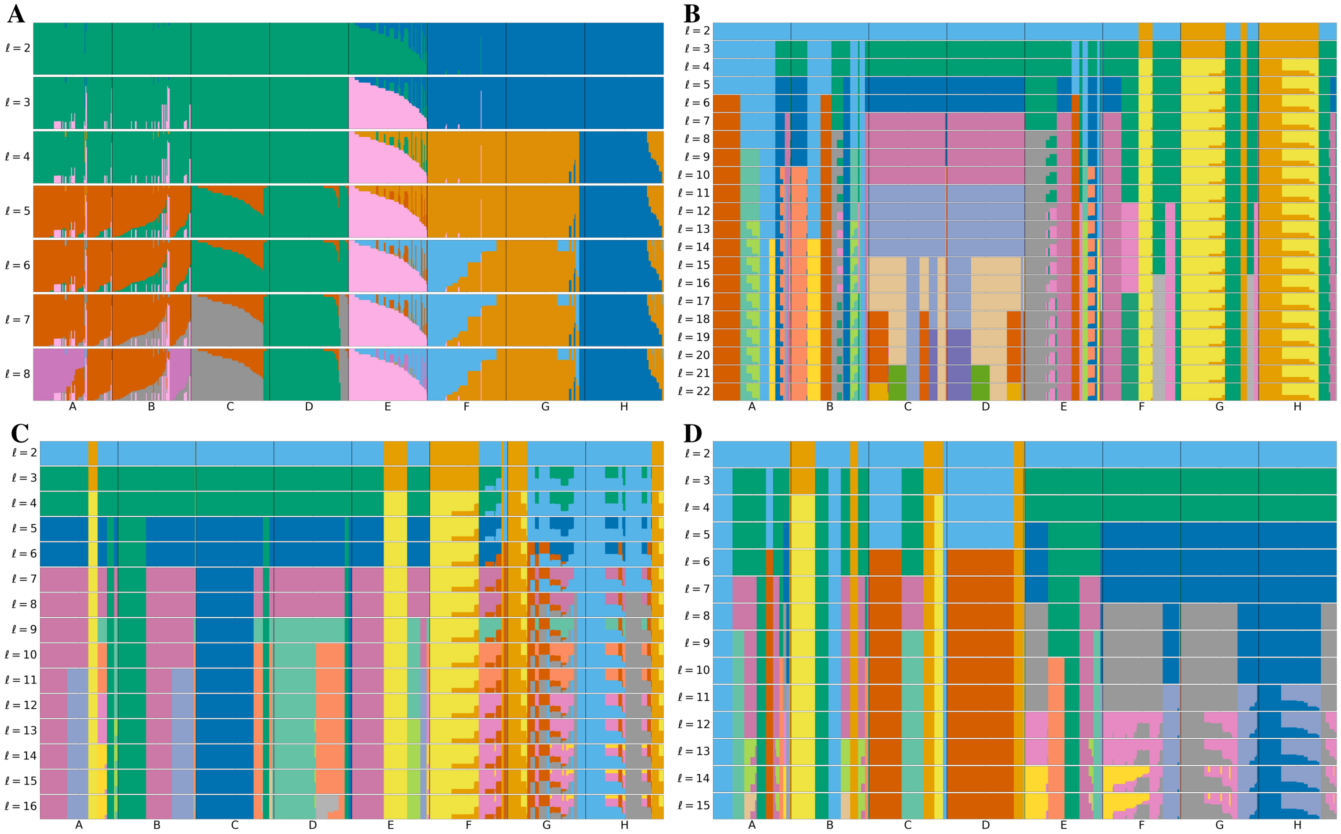


**Supplemental Fig. S15. Random cost function on simulated data with migration.** This analysis shows the effects of assigning random costs to each SNP/cut using the simulated data with migration added between populations A to H (Fig. 5 A). Panel **A** shows tangleGen with the FST-based cost function to facilitate comparison. Panels **B**, **C**, and **D** show tangleGen with random cost assignments using different seeds. The individuals are sorted within the populations based on the soft cluster proportions to achieve a block structure in the plots. Each subplot corresponds to a level $\ell$ in the tangles tree, where the different levels result from splits in the tangles tree. The agreement parameter is set to $a = 30$ in all panels and all external branches supported by only one SNP are pruned. In this analysis, the populations are less well differentiated compared to Fig. S14, resulting in significantly fewer SNPs with discriminatory power. Without a reasonable cost function, both population splits and the order of splits are meaningless and do not reflect the underlying population structure or migration patterns. In contrast to Fig. S14, tangleGen now mostly subdivides individual populations. Additionally, the depth of the inferred hierarchy varies depending on the SNPs used first and has increased significantly. Furthermore, the inference is based on even fewer SNPs per level than in Fig. S14.

#### Supplemental Note 13: Effect of the Penalizing Factor $c_2$

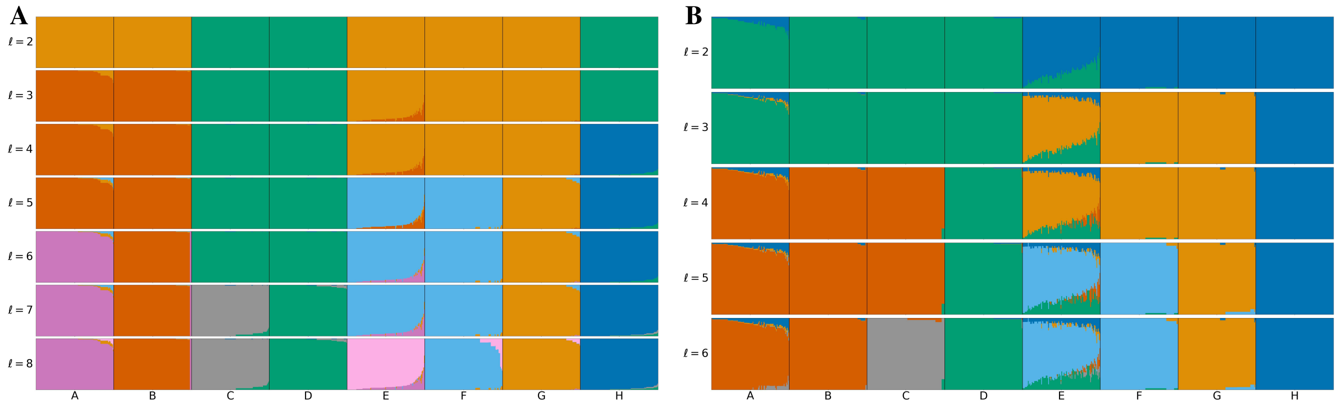


**Supplemental Fig. S16.** The FST-based cost function contains the penalty factor $c_2$, which slightly favors cuts that divide individuals into equally sized groups. This preference is meaningful for a hierarchical method but might reduce sensitivity to underrepresented, isolated populations. The simulated scenario in Fig. 4 A is suitable for evaluating this question, as the left subtree is balanced while the right subtree is unbalanced. The data used for panels A and B is identical; only the penalizing factor $c_2$ differs. **A: The penalty factor $c_2$ is deactivated.** The top split is now based on only one SNP, which is usually penalized by $c_2$ to be considered at a later stage. Other splits in the right, unbalanced subtree remain unchanged. **B: Increased penalty factor $c_2$.** To examine the other extreme, we increased the hyperparameter $b$ in the cost function, Eq. 2 in Methods, from $b = 0.05$ to $b = 0.5$. The root split works well, and H is still separated due to its high differentiation. However, ABC is separated at $\ell = 4$ instead of AB because separating ABC from DEFGH is more favorable than separating AB from CDEFGH. Furthermore, EF and AB can no longer be separated here. Panels A and B show that using the penalty factor $c_2$ is beneficial. The plots suggest that our choice of hyperparameter $b$ is appropriate, which is further supported by the hyperparameter analysis in Supplemental Note 11. Additionally, tangleGen can effectively detect underrepresented populations (such as H) as long as there are SNPs distinguishing these populations from the rest. If no such SNPs exist, detection would likely be challenging even without the $c_2$.
